# Dynamic sampling bias and overdispersion induced by skewed offspring distributions

**DOI:** 10.1101/2021.03.09.434663

**Authors:** Takashi Okada, Oskar Hallatschek

## Abstract

Natural populations often show enhanced genetic drift consistent with a strong skew in their offspring number distribution. The skew arises because the variability of family sizes is either inherently strong or amplified by population expansions, leading to so-called ‘jackpot’ events. The resulting allele frequency fluctuations are large and, therefore, challenge standard models of population genetics, which assume sufficiently narrow offspring distributions. While the neutral dynamics backward in time can be readily analyzed using coalescent approaches, we still know little about the effect of broad offspring distributions on the dynamics forward in time, especially with selection. Here, we employ an exact asymptotic analysis combined with a scaling hypothesis to demonstrate that over-dispersed frequency trajectories emerge from the competition of conventional forces, such as selection or mutations, with an emerging time-dependent sampling bias against the minor allele. The sampling bias arises from the characteristic time-dependence of the largest sampled family size within each allelic type. Using this insight, we establish simple scaling relations for allele frequency fluctuations, fixation probabilities, extinction times, and the site frequency spectra that arise when offspring numbers are distributed according to a power law *n*^−(1+*α*)^. To demonstrate that this coarse-grained model captures a wide variety of non-equilibrium dynamics, we validate our results in traveling waves, where the phenomenon of ‘gene surfing’ can produce any exponent 1 < *α* < 2. We argue that the concept of a dynamic sampling bias is useful generally to develop both intuition and statistical tests for the unusual dynamics of populations with skewed offspring distributions, which can confound commonly used tests for selection or demographic history.

## I. INTRODUCTION

Interpreting the genetic differences between and within populations we observe today requires a robust understanding of how allele frequencies change over time. Most theoretical and statistical advancements have been based on the Wright-Fisher model [1, 2], which has shaped the intuition of generations of population geneticists for how evolutionary dynamics works [3]. The Wright-Fisher model assumes that the genetic makeup of a generation results from resampling the gene pool of the previous generation, whereby biases are introduced to account for most relevant evolutionary forces, such as selection, migration, or variable population sizes. For large populations, the resulting dynamics can be approximated by a biased diffusion process, which simplifies the statistical modeling of the genetic diversity. More importantly, Wright-Fisher diffusion is the limiting allele frequency process of a wide variety of microscopic models, as long as they satisfy seemingly mild assumptions (see below). This flexibility has made Wright-Fisher diffusion the standard model of choice to infer the demographic history of a species, loci of selection or the strength of polygenic selection [4–9].

Despite its versatility, Wright-Fisher diffusion can be a poor approximation when the population dynamics is driven by rare but strong number fluctuations. It is increasingly recognized that number fluctuations can be inflated for very different reasons. First, the considered species may have a broad offspring distribution, which occurs for marine species and plants with a Type III survivorship curve [10, 11] as well as viruses and fungi (reviewed in [12]). Broad offspring distributions also arise in infectious disease, when relatively few super-spreaders are responsible for the majority of the disease transmissions [13]. In the recent SARS-CoV-2 pandemic, for example, a strongly skewed offspring distributions were consistently inferred from both contact tracing data and infection cluster size distributions [14, 15]. Understanding allele frequency trajectories in these systems is extremely challenging, as statistical inference based on the Wright-Fisher model is often misleading (see e.g. [16]).

A second mechanism for strong number fluctuations are so-called jackpot events, which can occur in any species no matter the actual offspring distribution. Jackpot events are population bottlenecks that arise when the earliest, the most fit or the most advanced individuals have an unusual large number of descendants. Temporal jackpot events (“earliest”) were first discovered by Luria and Delbrück [17] and studied as a signal of spontaneous mutations in an expanding population. They observed that a phage resistant mutant clone can grow exceptionally large if the resistance mutation by chance occurs early in an expansion event. Despite being rare, these jackpot events are easily detectable in large populations because they strongly inflate the variance of the expected number of mutants and lead to power-law descendant distributions.

The very same descendant distribution arises in models of rampant adaptation and of background selection. In these models, mutations generate jackpot events when they arise within the few fittest individuals [18]. Jackpot events also arise in range expansions, where the most advanced individuals in the front of the population have a good chance to leave many descendants over the next few generations. This phenomenon of gene surfing can produce a wide range of scale-free descendant distributions [19–22].

To account for skewed offspring distributions, a number of theoretical studies have been conducted in the context of the coalescent framework. Based on this backward-in-time, a striking feature of broad offspring distributions is the simultaneous merging of multiple lineages. One of the most widely studied models is the beta-coalescent [23], which is a subclass of the Λ-coalescent and corresponds to the population dynamics with a power-law offspring number distribution ∝*u*^−1+*α*^. The case *α* = 1, called Bolthausen-Sznitman coalescent [24], has been shown to be the limiting coalescent in models of so-called “pulled” traveling waves, which describe the most basic scenarios of range expansions [25] and of rampant adaptation [18, 26–28]. Moreover, so-called “semi-pushed” traveling waves that contain some level of co-operativity, induced e.g. by an Allee effect, generate power-law offspring distributions with 1 < *α* < 2 [21], indicating that their coalescent is intermediate between the Bolthausen-Sznitman and Kingman coalescents.

The tractability of coalescent approaches make it particularly useful for inferring demographic histories and detecting outlier behaviors [29–31]. However, as it is notoriously difficult to integrate selection in coalescent frameworks, there is also a strong need for forward-in-time approaches that capture the competition between genetic drift and selection. While for *α* ≥ 2, the limiting allele frequency dynamics is given by the well-understood Wright-Fisher process, much less is known for the case *α* < 2. This is unfortunate because, as mentioned above, any exponent 1 ≤ *α* ≤ 2 can arise dynamically.

Recently, the forward-dynamics of the special case *α* = 1 was studied by one of the authors [32], finding that an emergent sampling bias generates strong deviations from Wright-Fisher dynamics. The sampling bias arises because, in each generation, an allele with high frequency can sample more often and, hence, deeper into the tail of the offspring distribution than an allele with small frequency. The major allele of a biallelic site, therefore, has with high probability a greater number of offspring per individual than the minority type. This sampling bias acts like a selective advantage of the major allele, but its average effect is compensated by rare frequency hikes of the minor allele so that the expected change in frequency only changes in the presence of genuine selection.

Here, we focus on the understudied case 1 < *α* < 2 intermediate between the known cases of *α* ≥ 2, corresponding to Wright-Fisher diffusion, and *α* = 1 described by jumps and sampling bias but vanishing diffusion. Similarly to the *α* = 1 borderline case, we find that a minorallele-suppressing sampling bias arises but that it is fading over time as the offspring distributions are sampled more and more thoroughly. This time-dependent sampling bias determines the scaling of the fixation probability, extinction time, stationary distribution, and site frequency spectrum. The combination of jumps and bias generates a so-called Levy-flight which controls the variability of allele frequency trajectories, for instance between unlinked genes or between populations. The flexibility of our model should enable to fit wide range cases that deviate from Wright-Fisher diffusion.

## II. SAMPLING ALLELE FREQUENCIES ACROSS GENERATIONS

To study the impact of broad offspring numbers, we consider an idealized, panmictic, haploid population of constant size *N* that produces non-overlapping generations in the following way. First, we associate with each individual *i* a “reproductive value” [1, 33] *U_i_*, which represents its *expected* contribution to the population of the next generation. The random numbers *U_i_* are drawn from a specified distribution *P_U_*. In a second step, we sample each individual in proportion to its reproductive value until we have obtained *N* new individuals representing the next generation.

Our model belongs to the general class of Cannings models [34]. The Wright-Fisher model is obtained if we choose *P_U_* to be a Dirac delta function, such that all individuals have the same reproductive value.

We focus most of our analysis on the dynamics of two mutually exclusive alleles, a wild type and a mutant allele. The dynamics of the two alleles is captured by the time-dependent frequency *X*(*t*) (0 ≤ *X* ≤ 1) of mutants. The wild type frequency is given by 1 − *X*(*t*). The total reproductive values *M* and *W* of the mutant population and the the wild type population, respectively, are given by

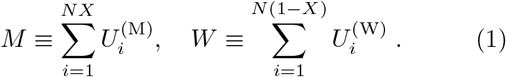

Here, 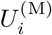 and 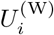 are the individual reproductive values of mutants and wild types and sampled from the distribution *P_U_*. The population at the next generation is generated by binomially sampling *N* individuals with success probability 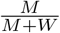. Mutations and selection are included as in the Wright-Fisher model. If the fitness of the mutant relative to the wild-type is 1 + *s*, where *s* is the selection coefficient, and the forward-and back-mutation rates are *μ*_1_ and *μ*_2_ respectively, then the success probability is given by 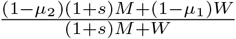.

For the offspring distribution *P_U_*, we consider a family of fat-tailed distributions, which asymptotically behave as 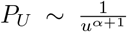 with *α* being a positive constant. To make our presentation concrete, we choose *P_U_* (*u*) = *α/u*^*α*+1^ (*u* ≥ 1), which is known as the Pareto distribution. In the large population size limit, the neutral allele-frequency dynamics is known to only depend on the asymptotic power law exponent *α* provided we measure time in units of the coalescence time [35].

## III. SIMULATION RESULTS

Our goal is to understand the asymptotic dynamics of our model for large *N*, where the frequency becomes continuous over time [36, 37] provided that *α* ≥ 1 [35]. We first present simulation results regarding relevant measures in the population genetics. Later, we provide a heuristic argument to explain them. Many separate observations (the fixation probability, extinction time, allele frequency fluctuations, stationary distribution, and site frequency spectrum) can be matched up with a unifying scaling picture.

Below, *t* and *τ* = *t/T_c_* denote a time in units of generations and one normalized by the characteristic (coalescent) timescale *T_c_*, respectively. *T_c_* depends on the population size and the exponent *α* as follows: *T_c_* = *N* when *α* > 2, *T_c_* = *N* / log *N* when *α* = 2, *T_c_* = *N*^*α*−1^ when 1 < *α* < 2, and by *T_c_* = log *N* when *α* = 1. These timescales were originally derived in the coalescent framework [35]. Later, we explain how they can be rationalized within the forward-in-time approach.

To understand the frequency dynamics when 1 ≤ *α* < 2, it is essential to distinguish between average and typical trajectories. As a proxy for typical trajectories, we use the median of the frequencies, denoted by *X*^med^(*τ*), throughout this paper.

### A. Neutral dynamics: typical trajectories and extinction time

First, we characterize the allele frequency dynamics in the absence of selection *s* = 0. In this neutral limit, the expected value of the allele frequency does not change over time, i.e., 〈*X*(*t*)〉 = *X*(0). Yet, despite the overall neutrality, a typical trajectory experiences a bias against the minority allele. This can be seen in Figure 1, where the mean and median are plotted across many realizations that start from the same frequency *X*(0) = 0.01. While the mean does not change over time, as required from neutrality, the median decays to zero in an *α*– dependent manner. By symmetry, the median increases towards fixation if the starting frequency is larger than 50%. Thus, the median experiences a bias against the minor allele. Note also that, when 1 < *α* < 2, the velocity of the median approaching extinction decreases as it approaches the extinction boundary (see the red curve in Figure 1). As we will show later, an uptick of the site frequency spectrum at the boundaries originates from this slowing.

**Figure 1.**
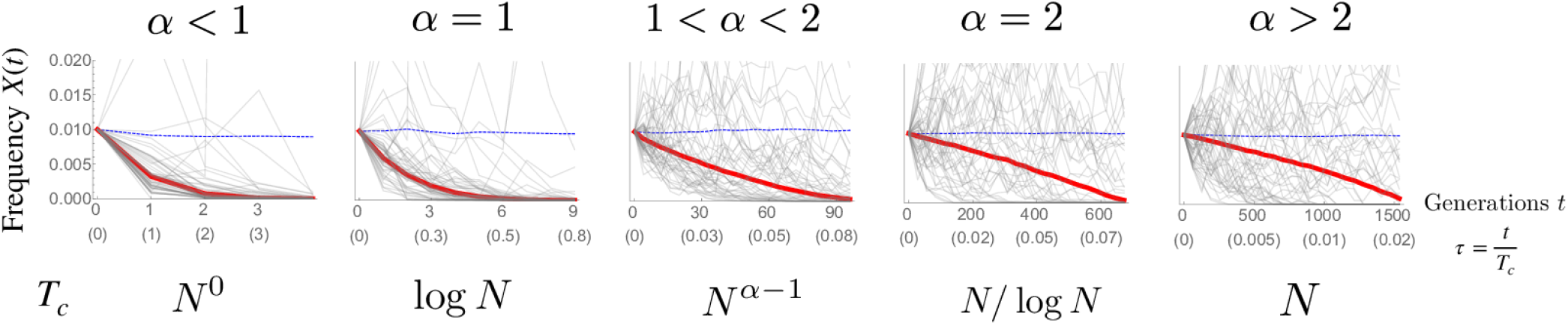
The mean (blue dashed curve) and median (red solid curve) of allele frequency trajectories for *α* = 0.8, 1, 1.5, 2, and 2.5. For each *α*, 10^4^ trajectories are generated with the initial frequency *x*_0_ = 0.01. For ease of viewing, only 50 trajectories are shown in gray in each panel. The time *t* in units of generations and the one 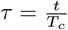 re-scaled by the coalescent timescale *T*_*c*_ are shown in the horizontal axes. The dependence of the coalescent time on the population size *N* is written below each panel. The population size is *N* = 10^5^.

Numerical simulations of the early part of trajectories show that time-dependent median displacement follows a simple power law,

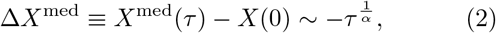

up to a frequency-independent prefactor. Figure 2 shows the numerical result for *α* = 1.5. The red curve represents the median of trajectories, which agrees well with 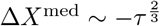.

**Figure 2.**
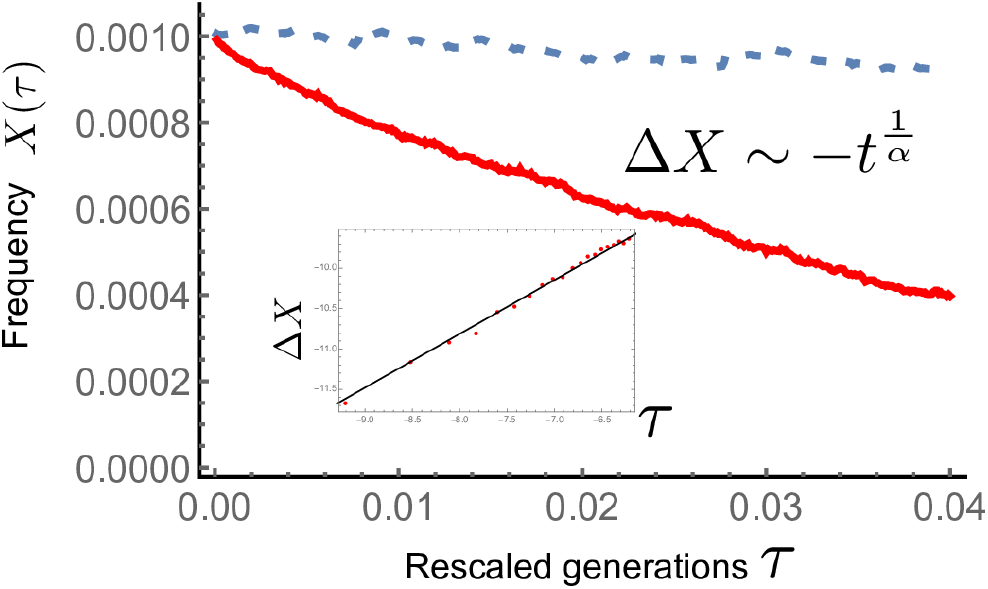
Trajectories of the median (red thick line) and the mean (blue dashed) of allele frequency when *α* = 1.5 and *N* = 10^8^. Inset: The trajectory (red points) of |Δ*X*^med^| = *X*(0) − *X*^med^(*τ*) is shown in log-log plot. The median agrees well with the expectation from the scaling argument 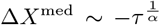 (black solid line).

Next we quantify the time to extinction, which turns out to be driven by the above minor-allele suppressing bias. Numerical results of the mean extinction time are consistent with

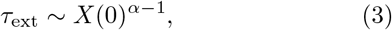

as shown in Figure 3. Hence, in units of the coalescence time, the mean extinction time *τ*_ext_ becomes larger as *α* decreases (namely, for a broader offspring distribution). Note, however, that if one measures time in units of generations, Equation 3 can be rewritten as *t*_ext_ = *τ*_ext_*T_c_* ~ (*NX*(0))^*α*−1^, which becomes smaller as *α* decreases since *NX*(0) ≥ 1.

**Figure 3.**
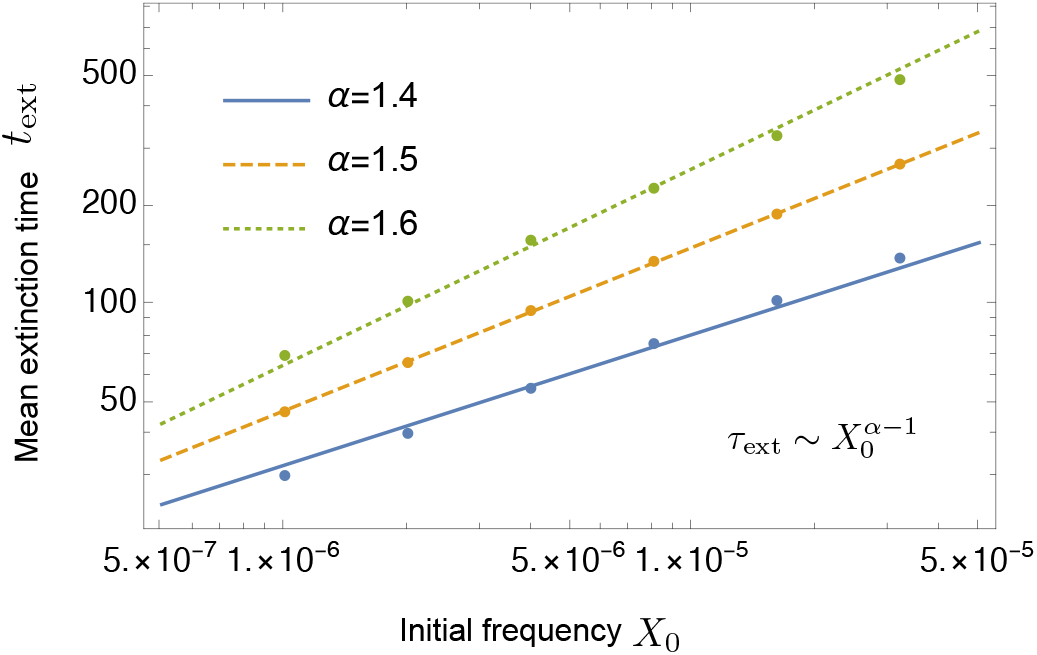
The mean extinction time *t*_ext_ (in units of generations) as a function of initial allele frequency *X*(0) is plotted for *α* = 1.4, 1.5, 1.6. Each of the straight lines has the slope *α* − 1. *t*_ext_ can be fitted well by Equation 3. The population size is *N* = 10^8^.

### B. Allele frequency fluctuations as a signature of broad offspring distributions

Next, we explore to what extent the spectrum of allele frequency fluctuations can provide a clue for identifying the exponent *α* of the offspring distribution. A deviation from the Wright-Fisher diffusion is most clearly revealed by measuring the median square displacement (median SD),

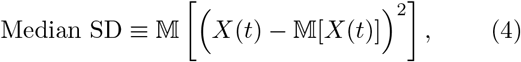

where 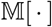 denotes taking the median (e.g. 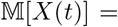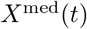. To measure the median SD, we simulate 1000 neutral allele frequency trajectories with initial condition *X*(0) = 0.5, for *α* = 1, 1.5 and the Wright-Fisher model (Figure 4A). As shown in Figure 4B, the median SD computed from this data set is consistent with the scaling,

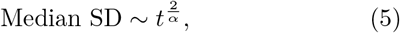

when *t/T_c_* ≪ 1. Noting 1 ≤ *α* < 2, this scaling means that typical fluctuations characterized by the median SD exhibit super-diffusion.

**Figure 4.**
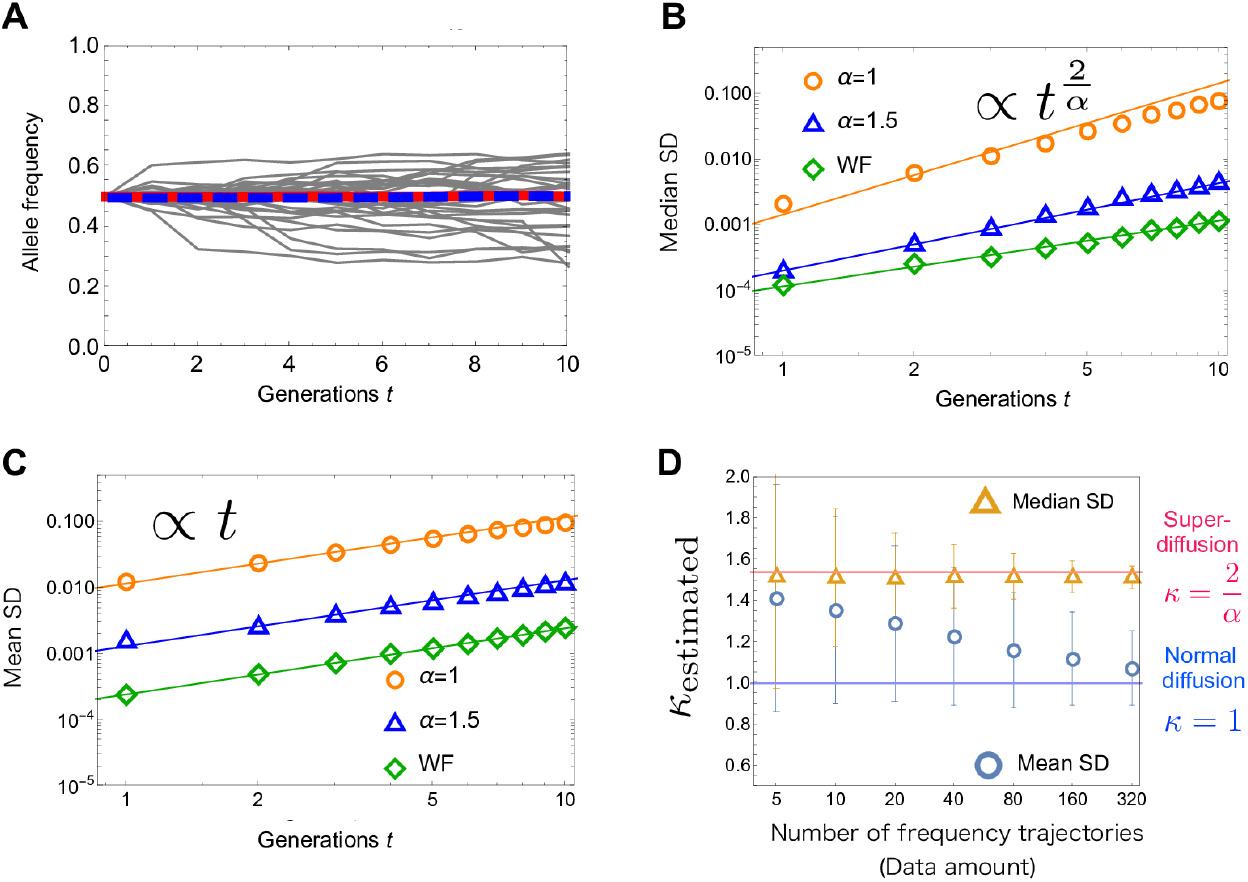
(**A**) Fluctuations of neutral allele frequencies when *α* = 1.5 and *N* = 10^5^. For *X*(0) = 0.5, the median *X*^med^(*t*) (red) is constant as well as the mean 〈*X*(*t*)〉 (blue). (**B**) The median square displacement computed from a data set of 1000 trajectories. For *α* = 1, 1.5 and the Wright-Fisher model, *N* = 10^8^, 10^4^, and 10^3^ are used respectively. The straight lines represent the scaling in Equation 5. For *α* = 1, the fitting after *t* ≳ 5 is not perfect, since *t/T*_*c*_ = *t/* ln *N* ≪ 1 is not satisfied. (**C**) The mean square displacement (mean SD) for different values of *α*. Solid lines represent linear scaling, which is expected for a regular diffusion process. (**D**) Data-size dependence of the estimated diffusion exponent *κ*_estimated_ for the mean SD (blue circle) and that for the median SD (orange triangle). See the main text for the detailed explanation. The horizontal lines show 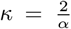 and *κ* = 1. The bars represent the standard deviations of *κ*_estimated_. *α* = 1.3 and *N* = 10^8^ are used.

Usually, allele frequency fluctuations are quantified by using the mean SD ≡ 〈(*X*(*t*) − *X*(0))^2^〉, rather than the median SD. For the Wright-Fisher diffusion, the distinction between these two measures is irrelevant since both of them increase linearly with time, except with differing prefactors. However, for 1 ≤ *α* < 2, the *α*-dependence in Equation 5 can be detected by measuring the median SD. As shown in Figure 4C, the mean SD (computed from a large data set) grows linearly in time even when *α* is less than 2, as if the underlying process was diffusive.

That the dynamics is not diffusive also impacts the mean SD, but somewhat subtly in that its value depends on the size of the data set (i.e., the number of frequency trajectories) used to measure it. This is because while rare large jumps contribute the mean SD in a large data set, these jumps are not observed in a small data set (with high probability). To demonstrate this data-size dependence, we prepare an ensemble of data sets, where each data set consists of a given number of allele-frequency trajectories. Then, for each data set, we measure the diffusion exponent *κ*, which is defined by

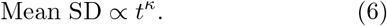

In Figure 4D, the ensemble-averaged exponent is shown by the blue circle. We can see that, as the data size increases, fluctuations characterized by the mean SD exhibit a crossover from super-diffusion 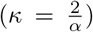 to normal diffusion (*κ* = 1). For the median SD, by contrast, we find that its diffusion exponent *κ* can be computed reliably without any significant dependence on the size of the data set (orange triangles in Figure 4D). For example, under the parameter setting in Figure 4D, given a date set of 320 trajectories, the diffusion exponent *κ*_estimated_ of the median SD falls within the interval [1.45, 1.57] with probability ~ 68%. This in turn predicts 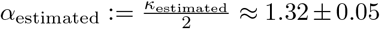, which is close to the actual value *α* = 1.3.

### C. Fixation probability

Next, we examine the effect of natural selection on the fixation probability of beneficial mutations. We consider a mutant with positive selective advantage *s* > 0 arising in a monoclonal population. The fixation probability *P*_fix_(*s*) of a single mutant depends on the parameter *α* of the offspring distribution. In the Wright-Fisher model (or equivalently, *α* ≥ 2), the fixation probability can be obtained using a diffusion approximation and is given by 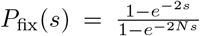, which becomes *P*_fix_ ≈ 2*s* when *Ns* ≫ 1 and *s* is small. When *α* = 1, an analytic result has been recently obtained in [32], which can be approximated as 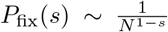. For the intermediate case, 1 < *α* < 2, we find that the fixation probability is given by

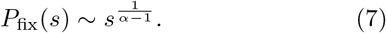

See Figure 5 for the numerical results. Note that since 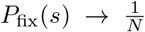 in the neutral limit independently of *α*, these results hold for sufficiently strong selection, 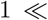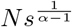.

**Figure 5.**
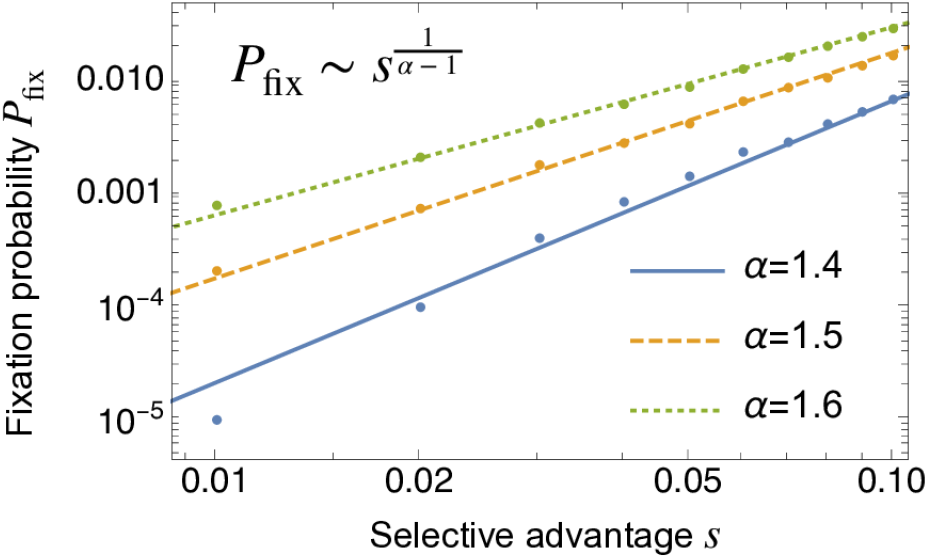
The fixation probability *P*_fix_ as a function of selective advantage *s*. The lines are the expectations from the scaling argument in Equation 42. The population size is *N* = 10^8^.

As Equation 7 shows, for a fixed population size and selective advantage, the fixation probability becomes smaller as *α* decreases. Intuitively, this is because, for smaller *α*, the success of fixation in catching a ride on a jackpot event depends more strongly on luck than on fitness differences.

### D. Site frequency spectrum

The site frequency spectrum (SFS) is often used as a convenient summary of the genetic diversity within a population. Theoretically, the SFS is defined in the infinite alleles model [38] as the density *f*_SFS_(*x*) of neutral derived alleles in the population (namely, *f*_SFS_(*x*)*dx* is the number of derived alleles in the frequency window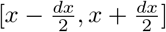.

Figure 6 shows numerical plots of the SFS for *α* = 1, 1.5, and the Wright-Fisher model. In the standard Wright-Fisher model, the SFS is proportional to 1*/x*, which decreases monotonically as *x* increases. By contrast, when offspring numbers are broadly distributed (when *α* < 2), the SFS is non-monotonic with a somewhat surprising uptick towards the fixation boundary. When *α* = 1, the analytic understandings of asymptotic behaviors near both boundaries are well-established: *f*_SFS_(*x*) is proportional to 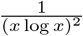 near *x* ~ 0 and 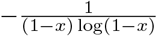 near *x* ~ 1, respectively [18, 27] (see also Appendix E).

**Figure 6.**
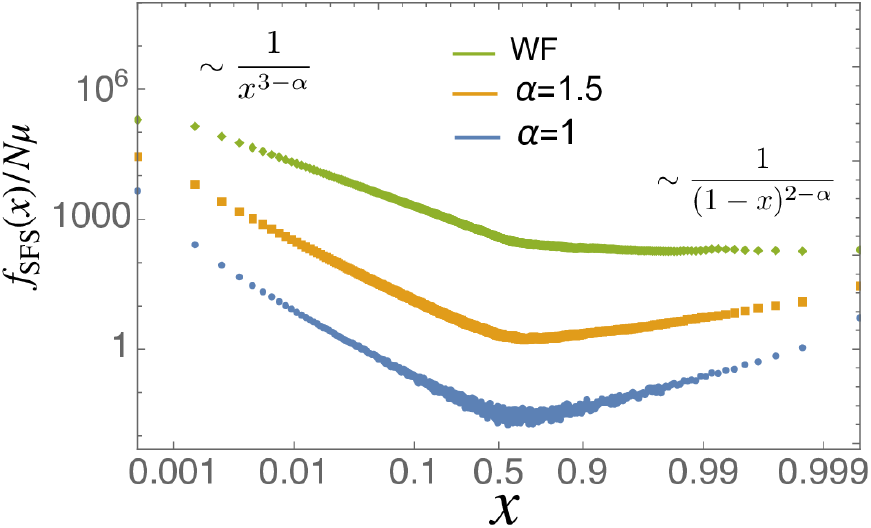
The site frequency spectrum for different values of *α* and fixed population size *N* = 10^5^. When 1 < *α* < 2, the rare-end spectrum and the frequent-end spectrum are 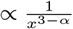 and 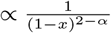, respectively (see also Figure 7).

For the intermediate case 1 < *α* < 2, the rare-end behavior of the SFS has been analytically studied. From a backward approach (the Λ-coalescent), the authors in [39] showed

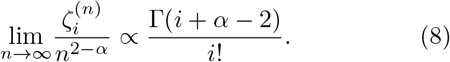

Here, *n* is a sample size and 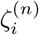 is the number of sites at which variants appear *i* times in the sample (see [39] for the proportionality constant of the right-hand side of Equation 8). By using Stirling’s approximation in Equation 8, we have

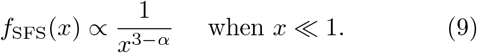

Equation 8, cannot be used for high-frequency variants, because the number of times the variants appear (*i* in Equation 8) is kept finite in taking the limit of the sample size *n*. To the best of our knowledge, a precise behavior at the high-frequency end for 1 < *α* < 2 has not been reported. As shown in Figure 7, we find that the asymptotic form of the uptick of *f*_SFS_(*x*) is given by

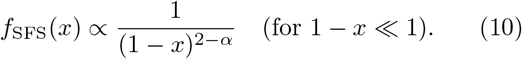

**Figure 7.**
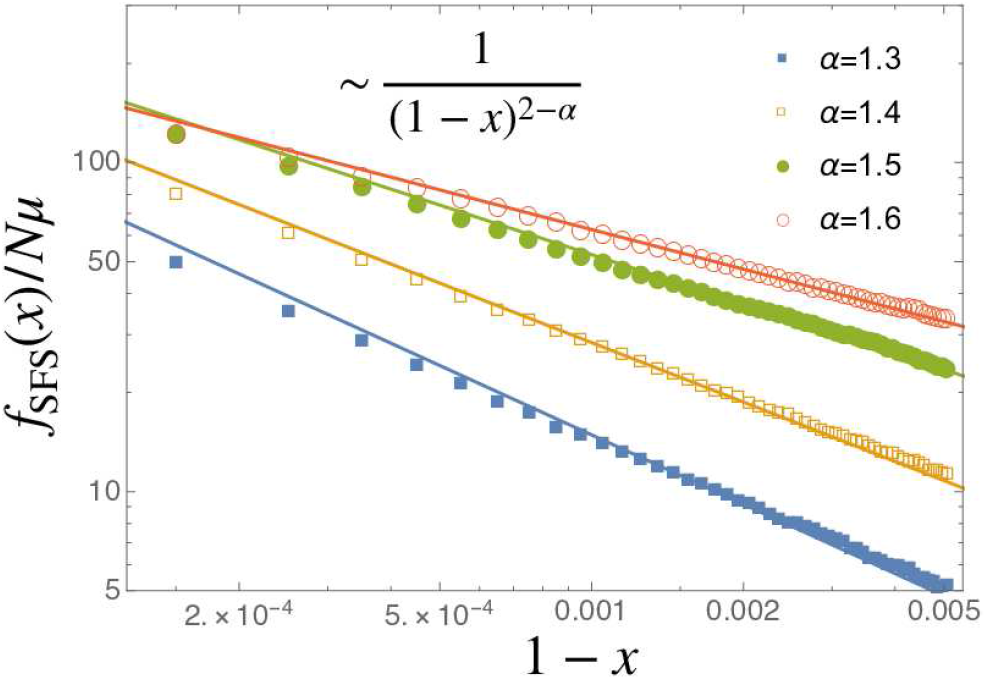
The SFS near *x* = 1 for *α* = 1.3, 1.4, 1.5, 1.6 (circle, squares). The horizontal axis is 1 − *x*. The solid lines are drawn assuming 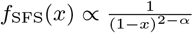. *N* = 10^6^ is used.

### E. Mutation-drift balance

A broad offspring distribution also affects the stationary distribution of allele frequency when mutations and genetic drift are balancing one another. For simplicity, we consider symmetric reversible mutations between two neutral allele types. We denote the scaled mutation rate (per unit time in the continuous description) as *θ* = *T_c_μ*, where *μ* denotes the mutation rate per generation. In the Wright-Fisher model, it is known that the stationary distribution is given by [36]

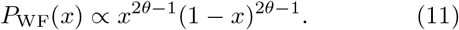

There is a critical value 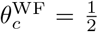: When 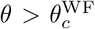, the distribution in Equation 11 has a single peak at the center 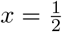; when 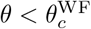, it has a U-shaped distribution, where the density is increasing monotonically from the center to the boundaries.

Figure 8A and B show the numerical results of the stationary distributions for the Wright-Fisher model and *α* = 1.5, respectively. When 1 ≤ *α* < 2, while a critical value of the mutation rate *θ_c_* exists as in the Wright-Fisher model, there is a qualitatively different feature: For a small mutation rate *θ < θ_c_*, the stationary distribution is not a U-shaped but an M-shaped distribution with two peaks near the boundaries. Note that the M-shaped distribution indicates a stochastic switching behavior, as illustrated in Figure 8D) (the blue curve). As shown in Figure 8D, the peak positions are approximately given up to prefactors bye

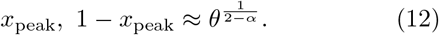

**Figure 8.**
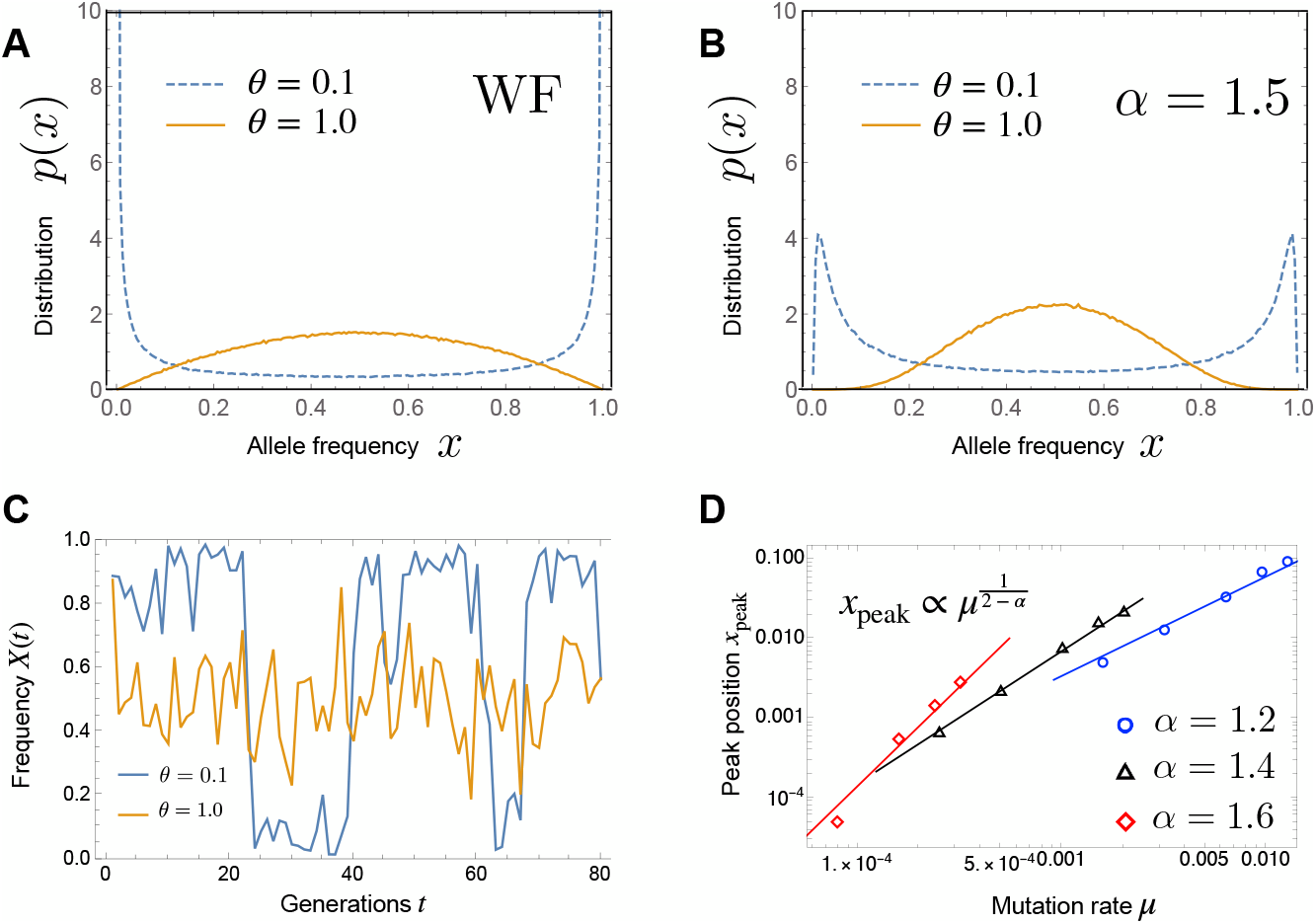
(**A**) Stationary distribution of the allele frequency in the Wright-Fisher model, when the mutation rate is small (*θ* = 0.1) and large (*θ* = 1.0). (**B**) Stationary distribution for *α* = 1.5, when the mutation rate is small (*θ* = 0.1) and large (*θ* = 1.0). (**C**) The time-series of the allele frequency in the case of *α* = 1.5, when the stationary distribution is bimodal (*θ* = 0.1) and unimodal (*θ* = 1.0). (**D**) The position of the peak near *x* = 0 of the stationary distribution versus the mutation rate *μ*. *N* = 10^4^ is used.

In Appendix H, we show that the M-shaped stationary distribution persists even in the presence of natural selection, provided that selection is weaker than the sampling bias at the peaks of the distribution.

A similar M-shaped distribution was observed for the EW process in [40], wherein moments of the stationary distribution were extensively studied. However, the origin of the M-shaped distribution remained unclear. Below, using scaling arguments, we explain why the bimodal distribution arises in our case.

## IV. ANALYTICAL ARGUMENTS

### A. Limiting process, transition density, and time-dependent effective bias

We now provide analytical arguments for the observations made in the simulations described in the first part of this paper. Our discussion starts with an exact but somewhat unwieldy description of the allele frequency dynamics. We then show how exact short-time and intermediate time asymptotics can be derived and used to rationalize the sampling bias and the scaling laws discovered above.

The allele frequency dynamics can be fully characterized by the transition probability density *w_N_* (*y|x*) that the mutant frequency changes from *x* to *y* in one generation. Since one generation consists of random offspring contributions to the seed pool and binomial sampling from the seed pool, we have

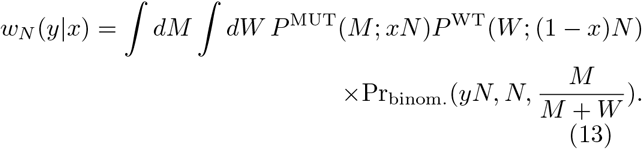

Here, *P* ^MUT^(*M*; *xN*) is the probability density that the sum of *xN* random mutant offspring numbers takes the value *M*, *P* ^WT^(*W*; (1 − *x*)*N*) is that for the wild type, and *Pr_binom_*. is the probability of getting *yN* successes in *N* trials with success probability 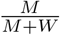. First, we will focus on the neutral case, for which *P* ^MUT^ and *P* ^WT^ are the same function, i.e., *P* ^MUT^(·) = *P* ^WT^(·).

While the resampling distribution *w_N_* may in general behave in complex ways, it has few options in the large *N* limit. These constraints emerge from two asymptotic simplifications. First, since *M* and *W* are the sums of many random variables, *P* ^MUT^ and *P* ^WT^ tend to stable distributions as described by the generalized central limit theorem [41, 42] (see also Appendix A for a brief description of the theorem). Second, the fluctuations associated with binomial sampling become negligible compared with those induced by offspring number contributions to the seed pool, provided that the offspring distribution is sufficiently broad, i.e., *α* ≤ 2. Thus, we can replace 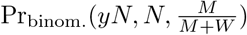 with a Dirac delta function, 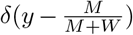. By using these facts and evaluating the integral in Equation 13 (see Appendix B for details), we obtain a simple analytical expression of *w_N_* (*y|x*), which is valid in the large *N* limit: When *α* = 1 [32],

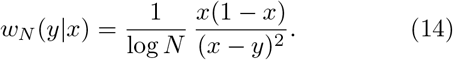

When 1 < *α* < 2,

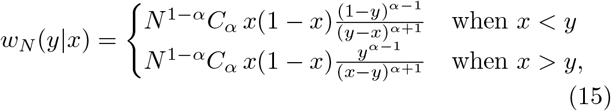

where 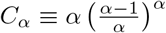.

To obtain the continuum description, we must appropriately scale the time *t* with the population size *N* [37]. The characteristic timescale (coalescent timescale) *T_c_* can be read from the dependence of the transition density on *N*. [32] showed that, when *α* = 1, the resulting limiting process is described by

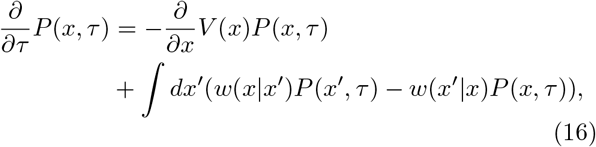

where the jump kernel *w*(*x*′|*x*) is given by

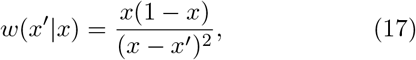

and the advection (bias) term *V*(*x*) is given by

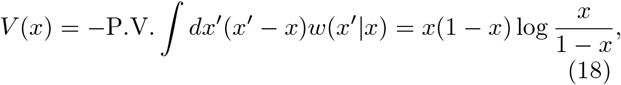

where P.V. denotes the Cauchy principal value. It is easy to check that Equation 18 satisfies the neutrality condition 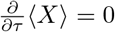.

To develop intuition, it is useful to interpret the different terms in Equation 16. First, *V* (*x*) has a form of frequency-dependent selection that enhances the major allele (with frequency > 50%) and suppresses the minor allele. The apparent fitness differences between the mutant and wild type is given by the log-ratio of their frequencies. Such a selection-like effect arises because the major allele can sample the offspring number from *P_U_* (*u*) more deeply than the minor allele (see [32]). Second, in spite of this apparent bias, the neutrality of the whole process is maintained due to rare large jumps, characterized by *w*(*y|x*). This also means that the neutrality does not hold if we focus on “typical” trajectories (see Figure 1). In fact, as we show in Appendix E, the median *x*_med_ of the mutant frequency, which is a proxy of “typical” trajectories, evolves according to

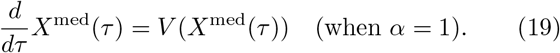

When 1 < α < 2, using the same reasoning as the derivation of Equation 16 and choosing 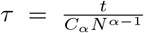 we can obtain the following differential Chapman-Kolmogorov equation,

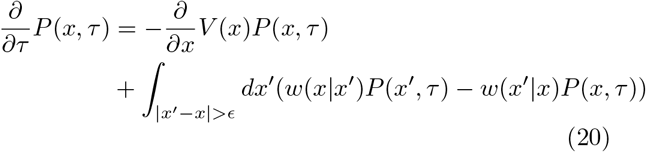

where

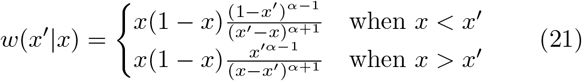

and

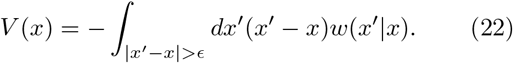

As in Equation 16, the advection term guarantees the neutrality of allele frequency. Equation 21 means that, when 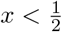, rightward jumps occur more frequently than leftward ones, and this tendency reverses when 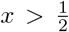. Noting the overall minus sign in Equation 22, this in turn means that *V*_eff_ is a bias against the minor allele (see Figure 1), as in the case of *α* = 1. We will later show that when *x* ≪ 1, the median trajectory is initially decaying like 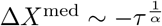 (Equation 2).

Note that, in the limit *ϵ* ⟶ 0, two divergencies arise in Equation 20, one in the integral for the advection velocity in Equation 22 and one in the jump integral in Equation 16. However, since both divergencies exactly cancel, the entire right-hand side of Equation 20 is well-defined. As shown in Appendix D, Equations 16 and 20 can also be derived as a dual of the Λ-Fleming-Viot process, namely as the adjoint operator of the backward generator (e.g., [43, 44]).

Although it is difficult to study Equation 20 analytically, it is possible to derive exact short-time and long-time asymptotics that, combined with scaling arguments, paint a fairly comprehensive picture of the ensuing statistical genetics.

### B. Short-time dynamics and fluctuations

First, we describe the transition density *P* (*x, τ|x*_0_*, τ* = 0) of Equation 20 for small times. When 1 < α < 2, the allele frequency changes due to the deterministic bias *V* (*x*) and random occurrence of jumps, sampled from the broad distribution in Equation 21. Since the number of jump events is enormous 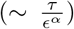 even for small *τ*, the generalized central limit theorem applies, and *X*(*τ*) is asymptotically distributed according to a stable distribution [41]. For a general stable distribution, its analytical expression is not available, and only its characteristic function can be expressed analytically. As we show in Appendix C, the random displacement Δ*X*(*τ*) = *X*(*τ*) − *x*_0_ can be expressed as

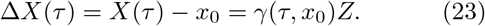

Here *Z* is sampled from the stable distribution *p*(*z*) whose characteristic function 〈*e^ikZ^*〉 ≡ ∫*dze^ikz^p*(*z*) is given by

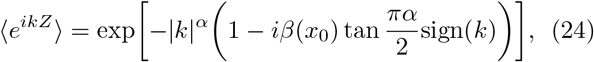

and the scale parameter *γ*(*τ, x*) and the skewness parameter *β*(*x*) are respectively given by

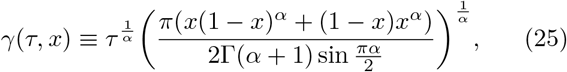

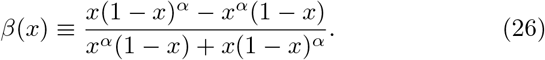

Note that statistical properties of *Z* are independent of *τ*, and Δ*X*(*τ*) depends on *τ* via the scale parameter *γ*(*τ, x*_0_). As shown in Figure 9A, for small times, the transition density *P* (*x, τ*|*x_0_, τ* = 0) computed from the stable distribution agrees precisely with numerical simulation results in the discrete-time model. Our result can be regarded as a counterpart of the Gaussian approximation often employed for Wright-Fisher diffusion (see [9] and the references therein).

**Figure 9.**
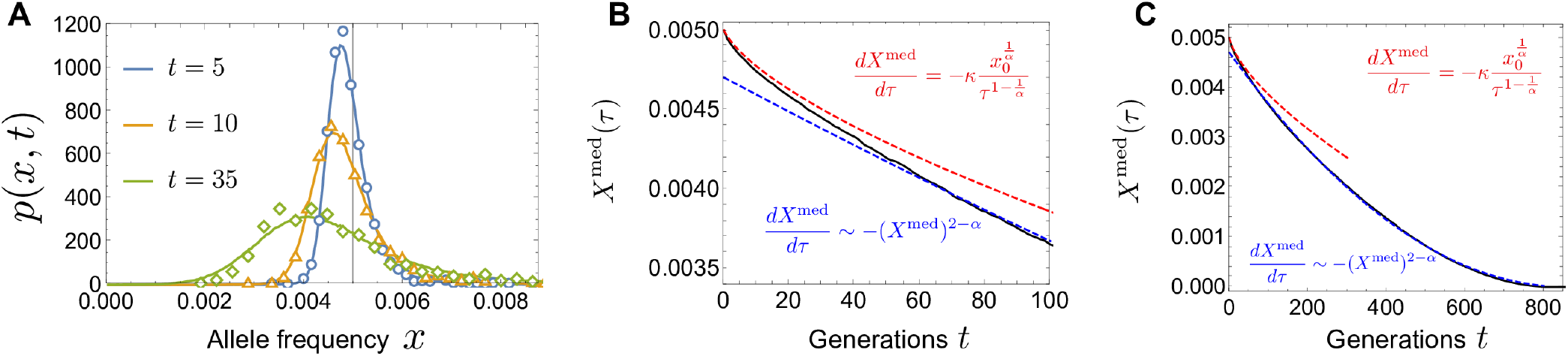
(A) The allele frequency distribution *p*(*x, t|x*0 = 0.005) at generation *t* = 5, 10, 35, for *α* = 1.5. The solid lines denote the short-time transition densities given by Equations 23 and 24, and the open markers denote those computed from 10000 allele frequency trajectories in the discrete-time model. (B) The initial dynamics of the median of the allele frequency (black). The red and blue lines denote the short-time solution in Equation 27 and the long-time solution in Equation 37, where constants of integration and the prefactor of Equation 38 are determined by fitting to the discrete-time model (black line) between 40 < *t* < 800. (C) The overall trajectory to extinction. The color scheme is the same as that in (B). In (A-C), *α* = 1.5, *N* = 10^7^, *x*_0_ = 0.005 are used.

Now, we study the mean and median of the allele frequency using the short-time expression. The mean does not change in time since 〈Δ*X*(*τ*)〉 = *γ*(*τ, x*_0_) 〈*Z*〉 = 0, which is consistent with the neutrality. On the other hand, the median changes as

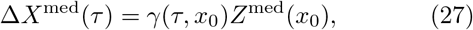

where *Z*^med^(*x*_0_) denotes the median of *Z*. *Z*^med^(*x*_0_) depends on *x*_0_ via *β*(*x*_0_) (see Equation 24), and *Z*^med^(*x*_0_) ≶ 0 for 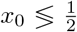. Equation 27 agrees with numerical simulations in the discrete-time model, while *X*(*τ*) is close to the initial frequency *x*_0_ (see the red and black curves in Figure 9 (B)).

The scaling property 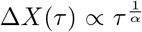 in Equation 2 immediately follows from Equation 27, since 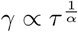. This scaling implies that there is a time-dependent bias driving the median of the allele frequency. Differentiating Equation 27 with respect to time gives

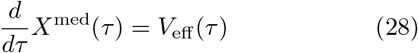

where the *effective time-dependent bias V*_eff_ (*τ*) is given by

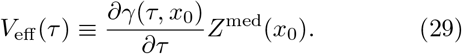

Near the boundaries *x* = 0 and *x* = 1, *V*_eff_ (*τ*) is approximately given by

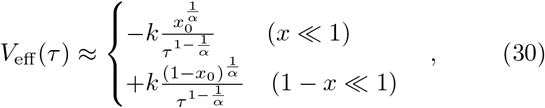

where 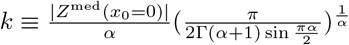 is a positive constant.

#### The advection term arises from a sampling bias

Intuitively, the time-dependent bias *V*_eff_ (*τ*) arises from a time-dependence of the largest sampled offspring number (Figure 10). To see this, consider a typical trajectory of the allele frequency starting from *x*. Up to a short time *τ*, only jumps from *x* to *y* ∈ [*y*_−_(*τ*)*, y*_+_(*τ*)] are likely to occur, where *y*_−_(*τ*) and *y*_+_(*τ*) can be estimated from

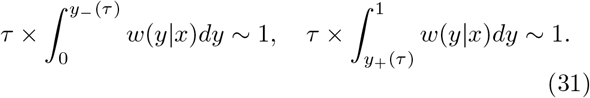

**Figure 10.**
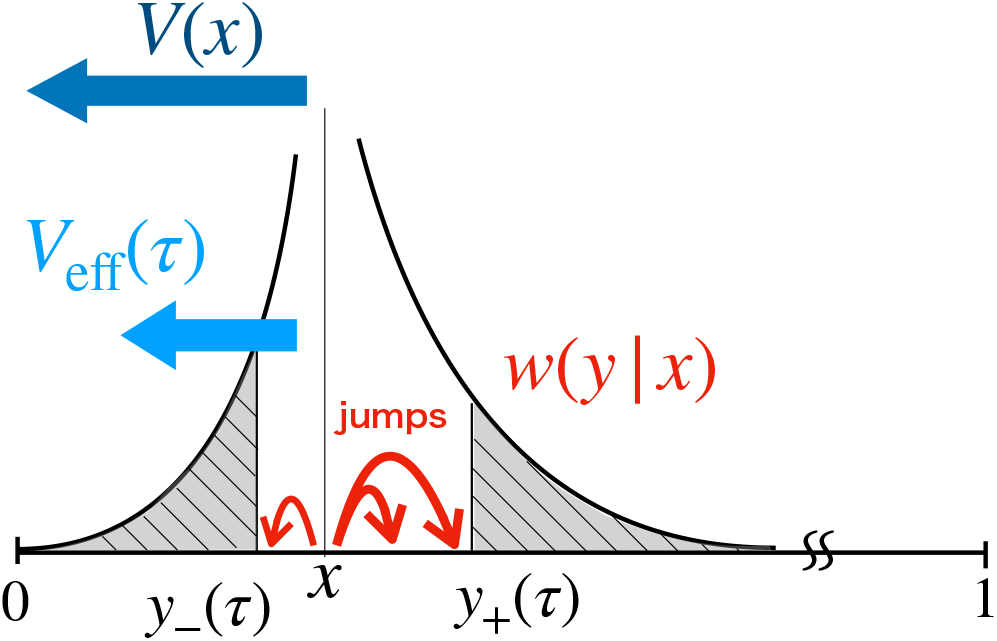
Schematic explanation of the effective time-dependent bias *V*_eff_(*τ*). The black curve shows the jump rate *w*(*y*|*x*) in Equation 21 when *x* ≪ 1. In a time *τ*, small jumps within the region [*y*−(*τ*)*, y*+(*τ*)] are likely to occur, offsetting a part of the original bias *V* (*x*). *V*_eff_(*τ*) is the residual part of the bias.

These conditions give

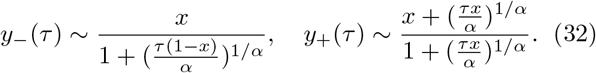

Because these small jumps cancel a part of the bias *V* (*x*) in Equation 22, the typical trajectory is then driven by the uncanceled residual part of the bias *V* (*x*),

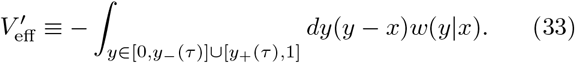

When *x*≪ 1, the dominant contribution to this integral is from *y* ≈ *y*_+_(*τ*). Using 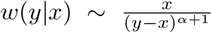 from the first line of Equation 21 and 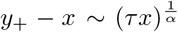 from Equation 32, the above integral can be evaluated as 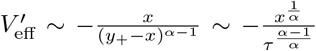, which agrees with *V*_eff_ in Equation 30 for *x* ≪ 1 (up to the factor *κ*). When 1 − *x* ≪ 1, the dominant contribution to 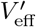 is from *y* ≈ *y*_−_(*τ*) and can be evaluated in a similar way, reproducing *V*_eff_ in Equation 30 for 1 − *x* ≪ 1.

#### Allele frequency fluctuations are inconsistent with Wright-Fisher diffusion

In the simulations, we found that, for 1 ≤ *α* < 2, allele frequency fluctuations are inconsistent with Wright-Fisher diffusion and characterized by super-diffusion with diffusion exponent 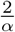 (see Equation 5). This finding is readily explained by the short-time asymptotic in Equation 23. Recalling statistical properties of *Z* are independent of *τ*, the median SD is given by

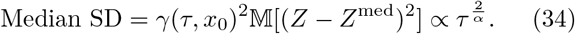

This scaling can also be justified heuristically by noting that, for 1 < α < 2, the square displacement is dominated by large jumps. During time *τ*, an allele frequency *X*(*τ*) around *x* typically jumps to *y*_±_ given in Equation 32. When *τ* ≪ 1, it is easy to see 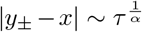 with *x*-dependent prefactors. Because the median SD is dominated by the largest displacements, it can be evaluated as

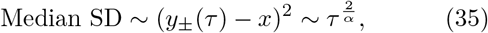

where 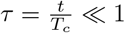 is assumed.

### C. Long-time dynamics and extinction time

Above, we saw that at short times, allele frequencies carry out an unconstrained Levy flight. This random search process, however, gets distorted as soon as the allele frequency starts to get in reach of one of the absorbing boundaries. Interestingly, the dynamics then enters a universal intermediate asymptotic regime that controls both the characteristic extinction time as well as establishment times and fixation probabilities.

To see this, let us consider the extinction dynamics of a trajectory starting from a small frequency *x*_0_ ≪ 1 (Figure 4). At short times, we can apply the short-time asymptotics in Equations 28, 30. We expect Equations 28, 30 to break down when the displacement Δ*X*^med^(*τ*) computed from Equation 28 becomes comparable to *x*_0_, which occurs at 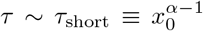. By taking a coarse-grained view, the rate of the frequency change in *τ*_short_ is roughly given by

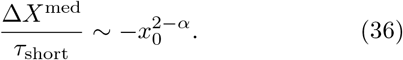

This suggests that, in a long timescale (*τ* ≳ *τ*_short_), the median frequency decreases as

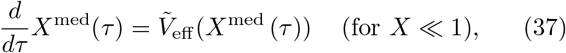

where, up to a prefactor, the frequency-dependent bias 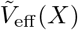 is given by

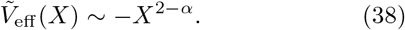

In Figure 9C, it is numerically shown that the long-time trajectory *X*^med^(*τ > τ*_short_) is consistent with Equation 37. By solving Equation 37, the median trajectory goes to extinction at 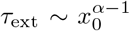 (Equation 3), in agreement with our simulations (Figure 4). Note that, for 1 − *x* ≪ 1, the bias in Equation 38 is replaced by 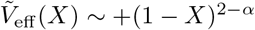.

Importantly, Equations 37 and 38 can also be rigorously justified from a scaling ansatz for the transition density. After some time, *P* (*x, τ|x*_0_) spreads broadly over the region *x* ≪ 1 with a peak at *x* = 0 (Figure 11A). As shown in Figure 11B, *P* (*x, τ*) is consistent with the following scaling ansatz;

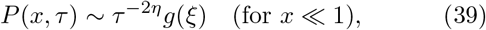

where 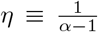 and *g*(*ξ*) is a function of 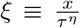. Up to an overall constant, *g*(*ξ*) can be determined analytically and expressed as an infinite series (see Appendix C). Note that the *τ* -dependent factor in Equation 39 is motivated from the fact that the extent over which the distribution spreads increases like *τ^η^*. Equation 39 implies that, conditional on establishment at *τ*, the median frequency increases as *X*^med^(*τ*)|_establish_ ~ *τ^η^*. Then, Equation 38 follows by evaluating the bias in Equation 30 at 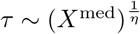 and at *X*^med^, instead of at *x*_0_.

**Figure 11.**
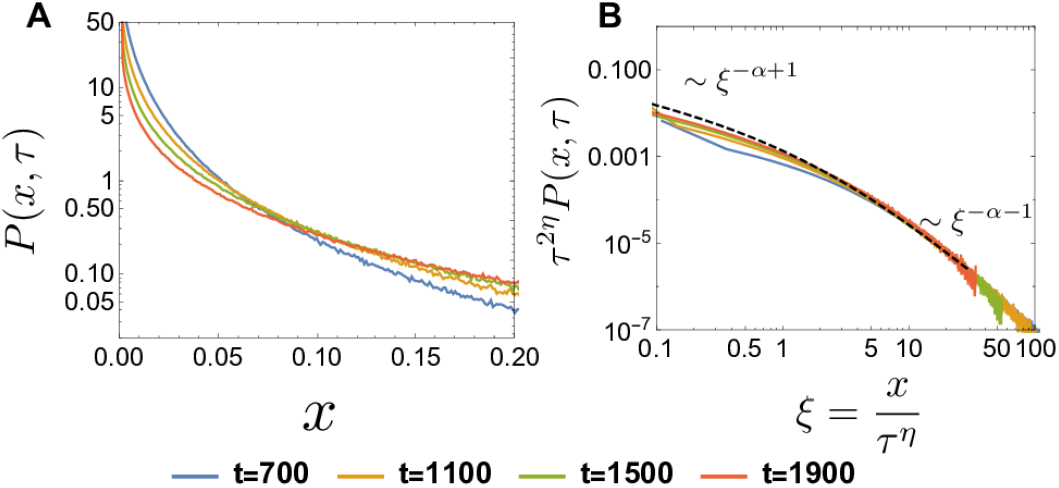
(A) Log plot of *P* (*x, τ|x*_0_ = 0.01) at generations *t* = 700, 1100, 1500, 1900 computed from the discrete-time model. *N* = 10^7^ and *α* = 1.5. (B) Log-Log plot of *τ*^2*η*^*P*(*x, τ|x*_0_ = 0.01) versus *ξ* = *x/τ*^2*η*^, where *η* = (*α* − 1)^−1^, at *t* = 700, 1100, 1500, 1900 (solid curves). The dashed curve represents the analytic result of *g*(*ξ*) (see Appendix C). The curves *τ*^2*η*^*P*(*x, τ|x*_0_ = 0.01) at the different time points collapse into *g*(*ξ*), supporting the scaling ansatz in Equation 39.

As a consistency check of the exponent *α* − 1 in Equation 3, we consider two solvable, extreme cases. First, in the limit *α* ⟶ 2, the dependence on *x*_0_ in Equation 3 becomes linear. In the Wright-Fisher model, the mean extinction time can be obtained analytically by solving the backward equation 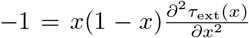 (see, for example, [45]). The solution is proportional to *x*_0_ with a logarithmic correction, *τ*_ext_ ≈ −*x*_0_ log *x*_0_. Second, when *α* ⟶ 1, the mean extinction time no longer depends on *x*_0_. We can obtain this explicitly, by solving Equation 19: Using *V*(*X*) ≃ *X* log *X* when *X* ≪ 1, the solution is given by 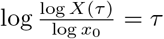. Therefore, if we approximately define the mean extinction time *τ*_ext_ as 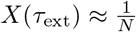, we obtain 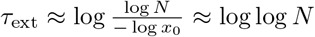, which is to leading order independent of *x*_0_ if *x*_0_ is taken to be of order one.

### D. Natural selection and fixation probability

One important advantage of the forward-time perspective is that we account for natural selection by introducing an appropriate bias favoring of the beneficial variant. Suppose that the mutant type has a selective advantage *s* > 0, such that the average offspring number of mutants is increased by a factor of 1 + *s* relative to the wild type. In time-rescaled Chapman-Kolmogorov equation, this adds the term *σx*(1 − *x*), where *σ* = *T_c_s*, into the advection *V* (*x*) of Equation 20.

The key observation underlying the argument below is that when *X* is sufficiently small, the selection force is negligible compared to the bias 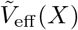 in Equation 38 because while the former is approximately linear in *X*, the latter is sublinear. If the frequency happens to grow and reach a certain value *X_c_*, the genuine selection begins to dominate over the bias, and the trajectory fixes with high probability (see Figure 12 for example trajectories and Figure 13)A). By using Equation 38, the crossover point *X_c_* can be estimated from balancing selection with the sampling bias,

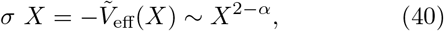

which gives

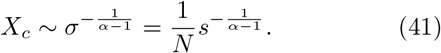

**Figure 12.**
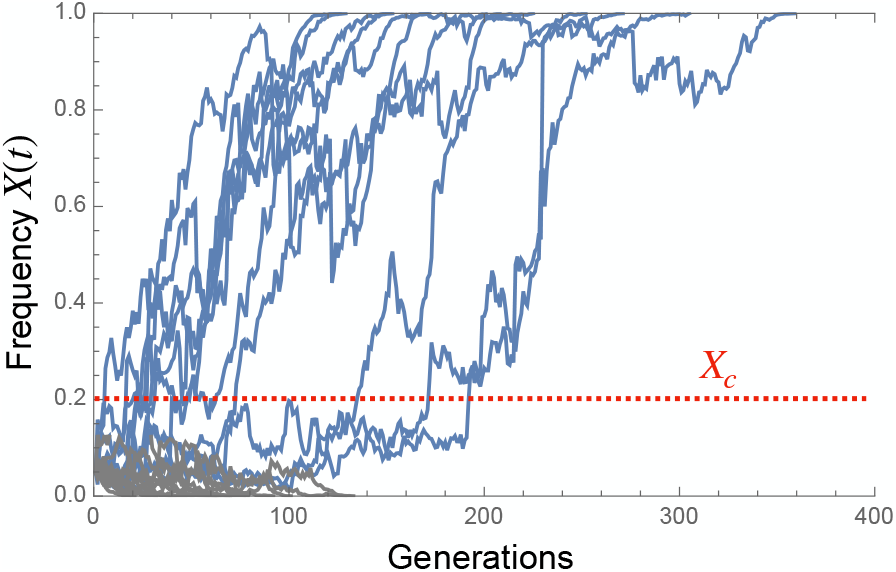
Example of trajectories of the frequency of the beneficial allele, starting from *x*_0_ = 0.05. *α* = 1.5*, s* = 0.03 and *N* = 5000. Fixed trajectories are colored in blue and extinct ones in gray. Here, the crossover point *X*_*c*_ can be estimated as *X*_*c*_ ~ 0.2 (, assuming that the proportional constant in Equation 41 is one). Once a trajectory reaches the crossover point, it becomes fixed in high probability.

**Figure 13.**
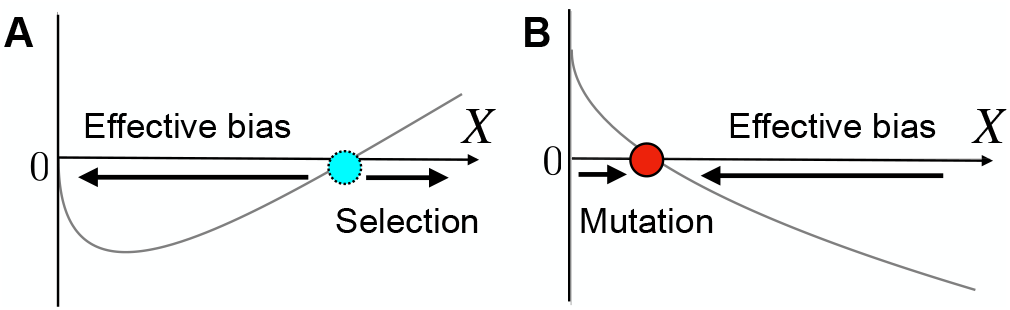
(A) The crossover from the effective bias to genuine selection. *V*(*X*) ~ − *CX*^2−*α*^ + *σX* is plotted, where *C* is a positive coefficient and *σ* > 0. Deterministically, an unstable point exists at *x* ~ *X_c_*. (B) The balance between the effective bias and mutation. *V*(*X*) ~ −*CX*^2−*α*^ + *θ* is plotted. Deterministically, a stable point exists at *X* ~ *θ*^1/(2−*α*)^.

The fixation probability *P*_fix_ can be estimated by using the neutral fixation probability in a population of size ≈*NX_c_*, because the dynamics are essentially neutral for *X ≪ X_c_*, and the trajectory grows almost deterministically for *X > X_c_*. Thus, the fixation probability is approximately given by

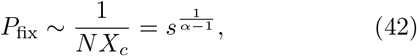

which is valid for 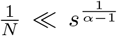. Equation 42 reproduces our simulation results in Figure 5 for 1 < α < 2 and, as *α* ⟶ 2, also reproduces the known result of the Wright-Fisher model, 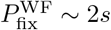 (up to a prefactor).

### E. Site frequency spectrum

By using the time-dependent effective bias, we can also estimate the behavior of the SFS *f*_SFS_(*x*) for frequent and rare variants. While the SFS is theoretically defined in the infinite alleles model, it can be computed from our biallelic framework ([46]): *f*_SFS_(*x*)Δ*x* is defined as the expected number of neutral derived alleles in the frequency interval 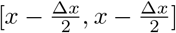 in a sampled population (here, the whole population). Because new mutations are assumed to arise uniformly in time, the SFS for unlinked neutral loci is given by the product of the total mutation rate *μN* and the mean sojourn time, namely, the average time an allele spends in the frequency interval 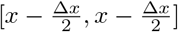 until fixation or extinction.

First, we consider the low-frequency end, *x* ≪ 1, of the SFS (see [47] for a similar argument). Since the SFS is proportional to the sojourn time, trajectories whose maximum frequencies are *x* or slightly larger than *x* dominantly contribute to the SFS *f*_SFS_(*x*) at *x*. Since these trajectories typically go extinct due to the bias, and we can roughly estimate their sojourn times at *x* as the inverse of “velocity”, 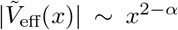 in Equation 38. Since the probability that a trajectory grows above a frequency *x* is roughly given by ~ 1/(*Nx*), the SFS is proportional to

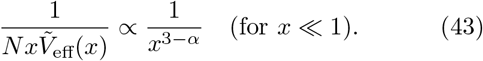

Similarly, for the high-frequency end of the SFS, only the trajectories that grow above *x* can contribute to *f*_SFS_(*x*). Typically, these trajectories go to fixation due to the bias 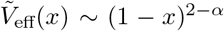. Therefore, the SFS is proportional to

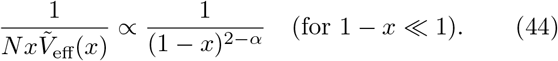

The effect of the genuine selection on the SFS can also be studied by using the effective bias. See Appendix F.

### F. Bimodality of stationary distribution

Now, we turn to explaining the bimodality observed at mutation-drift balance. We found that, when the mutation rates are small, the stationary allele frequency distribution is not a U-shaped, as expected from Wright-Fisher dynamics, but M-shaped, as shown in Figure 8. The M-shaped distribution arises from the balance between the mutational force and the effective bias (see Figure 13B). In the Chapman-Kolmogorov equation, the mutational force is given by

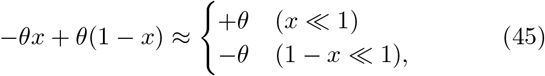

which pushes the frequency toward the center 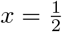. On the other hand, the effective bias, 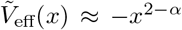 for *x* ≪ 1 and 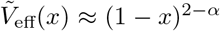 for 1 − *x* ≪ 1, pushes a trajectory toward the closer boundary. Therefore, the positions where these two forces balance are approximately given by

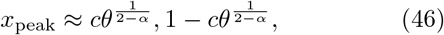

where *c* is a positive constant. If *θ* is sufficiently small, we can always find the balancing points. The presence of these two balancing points means that we can think of the allele frequency dynamics as a two-state system, essentially analogous to a super-diffusing particle in a double-well potential (see Figure 8C for a realization of trajectories). This explains the bimodal shape of the stationary distribution.

Finally, we remark that, even in the presence of natural selection, the balancing positions are still determined from the mutation-effective bias balance provided that *θ* ≪ 1: while the effective bias and the mutational term are sub-linear and constant respectively, the selection term *σx*(1 − *x*) is linear in *x* when *x* ≪ 1. Thus, when *θ* is sufficiently small, the magnitude of the selection term 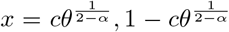 around is negligible, and the peak positions are given by Equation 46.

## V. DISCUSSION

In this study, we analyzed the effect of power law offspring distributions on the competition of two mutually exclusive alleles. Our main reason to consider such broad offspring distributions is that they often emerge in evolutionary scenarios that inflate the reproductive value [33] of a small set of founders. For example, range expansions blow up the descendant numbers of the most advanced individuals in the front of the population, an effect that has been called gene surfing [19]. Likewise, continual rampant adaptation boosts the descendant numbers of the most fit individuals. The resulting allele frequency dynamics becomes asymptotically similar to that of a population with scale-free offspring distributions.

In the case of narrow offspring distributions, which is predominant assumption in population genetics, it is usually an excellent approximation to describe the allele frequency dynamics by a biased diffusion process, which forms the basis of powerful inference frameworks [9]. If the offspring distribution is broad, however, allele frequency trajectories are disrupted by discontinuous jumps, resulting from so-called jackpot events - exceptionally large family sizes drawn by chance from the offspring distribution. Our goal was to find an analytical and intuitive framework within which we can understand the main features of these unusual dynamics.

We found that the main counter-intuitive features can be understood and well-approximated from a competition of selection and mutations with a time-dependent emergent sampling bias, *V*_eff_(*τ*). The sampling bias favors the major allele and arises, because the subpopulation carrying the major allele typically samples deeper into the tail of the offspring distribution than the minor allele fraction.

In the remainder, we first summarize the unusual population genetic patterns that can be explained by the action of these effective forces. We then discuss how broad offspring dynamics could be detected in natural populations and what its implications are for the dynamics of adaptation. Finally, we demonstrate that these dynamics are also ubiquitous in populations with narrow offspring distributions, when mutational jackpot are possible. Therefore, we believe our theoretical framework may be taken as a general null model for populations far from equilibrium.

### A. Unusual dynamics

We found that the sampling bias effectively acts like time- and frequency-dependent selection. In the absence of true selection, *V*_eff_(*x, τ*) drives the major allele to fixation, first rapidly and than gradually slowing down with time and proximity to fixation. The slowing down of the sampling bias near fixation also leads to an excess of high-frequency alleles, given continual influx of neutral mutations. This generates a high-frequency uptick in the site frequency spectrum, which is characteristic of the tail of the offspring distribution. In mutation-drift balance, the allele frequency distribution is M-shaped, in contrast to the U-shape expected from Wright-Fisher dynamics. The peaks reflect the balance of the mutational and sampling bias.

Non-neutral dynamics depends on whether the genuine selection force dominates over the sampling bias. The sampling bias tends to dominate near extinction or fixation, and wanes near 50% frequency. A de-novo beneficial allele will not be able to fix unless it overcomes, by chance, the switch-point frequency at which genuine selection becomes stronger than the sampling bias. Finally, fluctuations in typical trajectories are getting stronger over time. As a consequence, allele frequencies super-diffuse: fluctuations grow with time more rapidly than under regular Wright-Fisher diffusion.

### B. Detecting dynamics driven by broad offspring distributions

The time-dependent over-dispersion is most readily detected by plotting the median square displacement as a function of time (see Figure 4B). Testing deviations in this statistics are an attractive avenue for detecting deviations from Wright-Fisher diffusion because the signal is strong for intermediate allele frequencies, which can be accurately measured by population sequencing. By contrast, the time-dependent bias vanishes when an allele has 50% frequency. So, the detection of the sampling bias requires accurate time series data of low frequency variants, which is difficult to obtain given sequencing errors.

It is clear that a single super-diffusing but neutral allele would not abide by the diffusive Wright-Fisher null model and thus might be falsely considered as an allele under selection. But importantly, allele super-diffusion has an impact even on statistics that sum over many unlinked loci. This is significant for inference methods, for instance to detect polygenic selection, which argue that trait values follow a diffusion process, if not for an underlying Wright-Fisher dynamics of the allele frequencies then because they sum over many independent allele frequencies [6]. However, *α* < 2 dynamics breaks both of these arguments. In particular, sums of many unlinked loci tend to non-Gaussian distributions (so called alpha-stable distributions). Hence, for traditional inference methods based on Wright-Fisher diffusion or standard central limit theorem [9], an underlying super-diffusion process should be ruled out.

If time series are not available, broad offspring numbers can also be detected from the site frequency spectrum (SFS) [18]. A tail-tale sign of the sampling bias is a characteristic uptick at the high-frquency tail of the SFS, which is difficult to generate by demographic variation [18]. As we have shown, the shape of the uptick is characteristic of the tail of the offspring distribution (the parameter *α*).

### C. Implications for the dynamics of adaptation

We found that the fixation probabilities quite sensitively depends on the broadness *α* of the offspring distribution (Equation 42). Accordingly, the dynamics of adaptation, which ultimately depends on the fixation of beneficial variants, should change quantitatively. To estimate these modifications, we consider an asexual population of constant size *N* with a broad offspring distribution with 1 < α < 2, wherein beneficial mutations occur at the rate *μ*_B_. For low mutation rates, mutations sweep one after the other but when mutation rate are sufficiently high, multiple mutations occur and most mutations are outcompeted by fitter mutations. Such a situation is known as clonal interference.

We can study the effect of the exponent *α* on the adaptation dynamics quantitatively by repeating the argument in [48], wherein the variance of offspring numbers is assumed to be narrow. As discussed in Appendix G, clonal interference should occur if

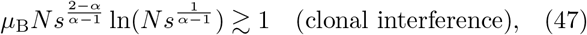

where *s* > 0 is the fitness effect of a mutation, which we assume to be constant. The rate *R* of adaptation is given by

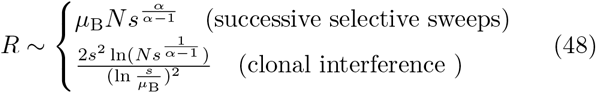

Note that the second line in Equation 48 reproduces Equation 5 of [48] in the limit *α* ⟶ 2. Thus, the rate of adaptation depends only weakly (logarithmically) on *α* in the clonal interference regime, even though the condition for clonal interference in Equation 47 depends on *α* quite sensitively,.

### D. Emergence of skewed offspring distributions in models of range expansions

Our study can be regarded as an analysis of the population genetics induced by power-law offspring distributions. The main reason to consider these scale-free offspring distributions is that they quite generally *emerge* in models of stochastic traveling waves [21]. Such models are ubiquitous in population genetics because they describe a wide range of evolutionary scenarios, including range expansions, rampant asexual and sexual adaptation as well as Muller’s ratchet [18, 21, 25–28]. Our analysis should therefore apply most directly to these evolutionary scenarios, which we now demonstrate using a simple model of a range expansion. We end by discussing the question of whether some of our results may also arise in scale-rich offspring distributions.

Ref. [21] argued that any exponent 1 ≤ *α* ≤ 2 can emerge in a simple model of range expansions that incorporates a tunable level of cooperativity between individuals (Figure 14A). The model can be described by a generalized stochastic Fisher-Kolmogorov equation

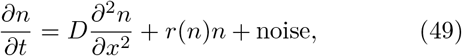

for the time-dependent population density *n*(*x, t*) at position *x* in a linear habitat and time *t*. The growth rate *r*(*n*) is assumed to be density-dependent, with

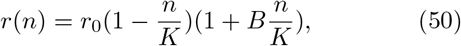

where the parameter *B* ≥ 0 accounts for co-operativity among individuals, which is also called an Allee effect. As discussed in [32], lineages in the region of the wave tip are diffusively mixed within the timescale 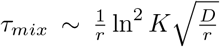. This implies that, in this microscopic model, resampling from an offspring distribution roughly occurs every *τ_mix_* generations. In [21, 22], it was argued that depending on the strength of the Allee effect, the offspring distributions corresponding to any of the three distinct classes of the beta coalescent process can arise; namely, the Bolthausen-Sznitman coalescent when *B* < 2, the beta coalescent with 1 < *α* < 2 when 2 < *B* < 4, and the Kingman coalescent when *B* > 4.

**Figure 14.**
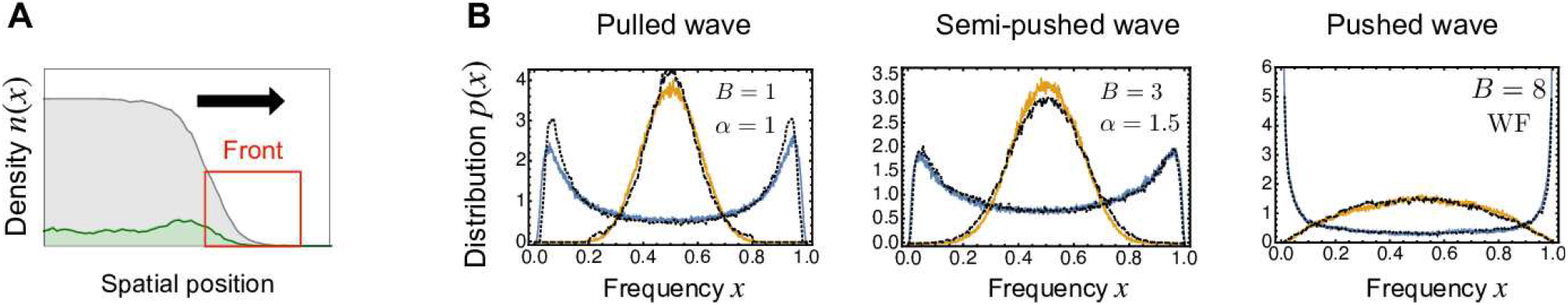
(**A**) The model of a range expanding population with two neutral alleles (green and gray). A broad offspring distribution arises dynamically in the front region. (**B**) Stationary distributions of the allele frequency when mutation rate *θ* is small (blue) and when *θ* is large (orange). The wiggling lines (blue/orange) are the numerical results in the traveling wave model, while the dotted lines (black) are those in the macroscopic model. The parameters of the Allee effect *B* are *B* = 1 (left), 3 (middle), and 8 (right). See Appendix I for the details of the implementation of the simulation and other parameter values.

To demonstrate clearly that our present study can serve as a macroscopic analysis of the traveling model, we introduce reversible mutations in the traveling wave model and measured the mutant frequency of the first 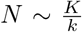 individuals from the edge of the front. Here, *k* is the spatial decay rate, i.e., 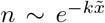 where 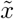 is the coordinate comoving with the expansion. This definition of the mutant frequency is reasonable because only the wave front has a skewed offspring distribution due to the founder effect. In Figure 14B, for *B* = 1 (left), 3 (middle), and 8 (right), the frequency distributions in the traveling wave model are shown when the mutation rate is small (orange jagged line) and when it is large (blue jagged line). The corresponding distributions in the macroscopic model are shown by black dotted lines. The stationary distributions in the traveling wave model agree well with those in the macroscopic model. Especially, the transition from the M-shaped or U-shaped distribution to the monomodal distribution is consistently reproduced in the traveling wave model. These results underscore the correspondence between the traveling wave with the Allee effect and the beta coalescent process.

The above-described correspondence suggests that the spatial area occupied by one allele type in a range expansion should behave statistically like the time-integral over the allele frequency in the Cannings model. In the context of adapting (non-spatial) populations, this quantity describes the total number of mutational opportunities of a mutant lineage [48–50]. As presented in Appendix J, the distribution of the time-integrated frequency exhibits a scaling behavior, that depends on the offspring distribution sensitively. While a full discussion is beyond the scope of this paper, we expect that the distribution of areas serves as a useful observable to distinguish different prototypes of traveling waves [21].

#### Broad offspring distributions with a scale

While scalefree offspring distributions often emerge over an intermediate time scale (*τ_mix_* in the above traveling wave model), there are also species that over single generations show broad offspring numbers and violate Wright-Fisher diffusion. For such species, it may be more natural to consider offspring distribution with a characteristic scale. In ‘sweepstake’ reproduction [11], a fixed and finite fraction of the population is replaced at every sweepstake event (specified by the parameter Ψ in [11]). Because Ψ sets a characteristic scale in offspring numbers, power law relationships for the median of allele frequencies as well as frequency fluctuations cannot be expected, which we confirm in Appendix K. Nevertheless, the qualitative features of a sampling bias can be recognized quite clearly for sweepstake reproduction as well.

Either type of model ultimately is an approximation to true offspring distributions, and it depends on the situation, which one to use. As we argued, the beta-coalescent along with the forward-in-time model described in this paper is the natural choice for range expansions, rapid adaptive process or other scenarios where the reproductive value of a chosen few are highly inflated.

## ACKNOWLEDGEMENTS

This work is in part supported by RIKEN iTHEMS Program. Research reported in this publication was supported by the National Institute of General Medical Sciences of the National Institutes of Health under award R01GM115851, a National Science Foundation CAREER Award (#1555330), a Simons Investigator award from the Simons Foundation (#327934), and JSPS KAKENHI (Grant Number JP19K03663). We express our sincere thanks to Benjamin H. Good, Daniel B. Weissman, Jiseon Min, Joao Ascensao, Michael M. Desai, and Stephen Martis for their helpful discussions and comments.

## Appendix A Generalized central limit theorem

Here, we briefly summarize the generalized central limit theorem [41, 42]. Suppose that each random number *u_i_* is sampled from the Pareto distribution 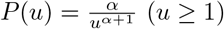) and consider the shifted and rescaled random variable *ζ*;

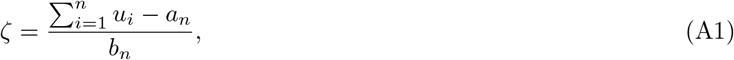

where *a_n_* and *b_n_* are

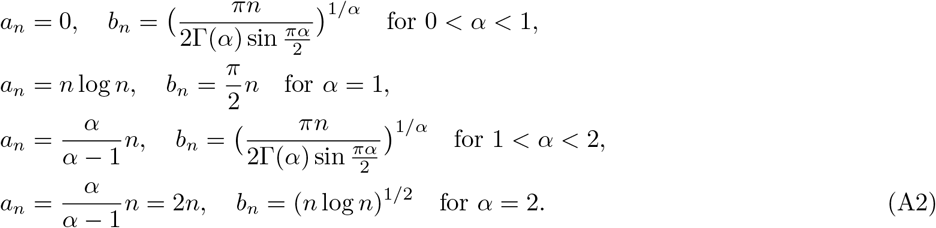

It is well-known that the distribution of *ζ* is well-approximated by the *α*-stable distribution, which we denote as *P_α_*(*ζ*). While an explicit expression of *P_α_*(*ζ*) is not available in general, the characteristic function is given by

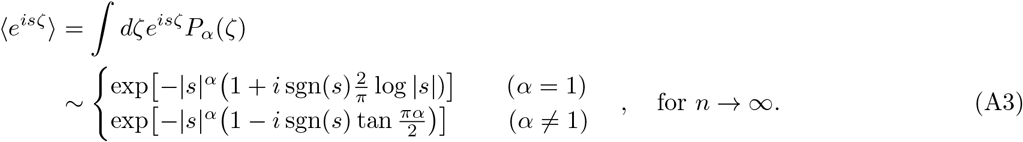

## Appendix B. The transition density of an allele frequency *w_N_*(*y*|*x*) and the asymptotic dynamics for lager *N*

Allele-frequency change in a generation is characterized by the transition density *w_N_*(*y|x*), which is the probability distribution of the allele frequency *y* at the next generation given the current allele frequency *x*. When *N* is lager, the asymptotic dynamics can be described by a time-continuous differential Chapman-Kolmogorov equation, which is defined by an advection velocity *V* (*x*), diffusion coefficient *D*(*x*), and jump kernel *w*(*y*|*x*) [37]. The triplet is obtained from the transition density *w_N_*(*y*|*x*) as follows:

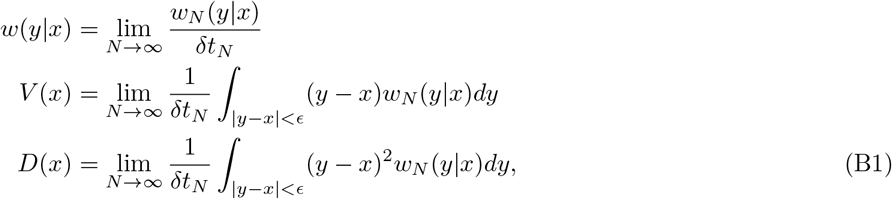

where *δt_N_* is an *N*-dependent timescale, corresponding to one generation measured in units of the coalescent timescale. In the following, we derive the transition density *w_N_*(*y|x*) and the asymptotic dynamicsfor general *α* by using a similar computational technique used in [32], wherein the case of *α* = 1 is studied extensively.

As mentioned in the main text, when *α* ≤ 2, the binomial sampling error is negligible for large *N* compared to the stochasticity coming from broad offspring number fluctuations, and we can replace the binomial distribution in Equation 13 of the main text with the Dirac delta function;

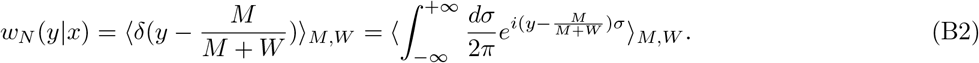

Here 〈·〉_*M,W*_ means the average over 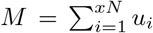 and 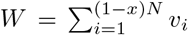. Using the variable 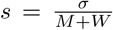, we can rewrite *w_N_* as

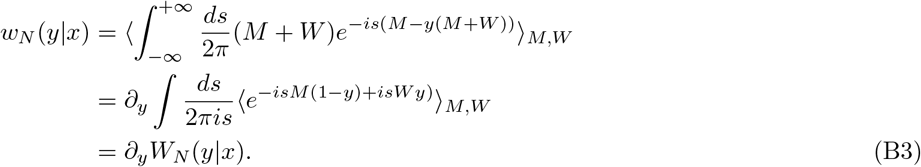

Here,

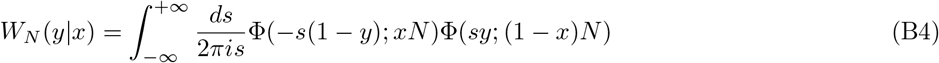

with

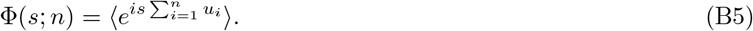

To use the properties of the *α*-stable distributions in Appendix A, we further rewrite *W_N_*(*y*|*x*) as follows:

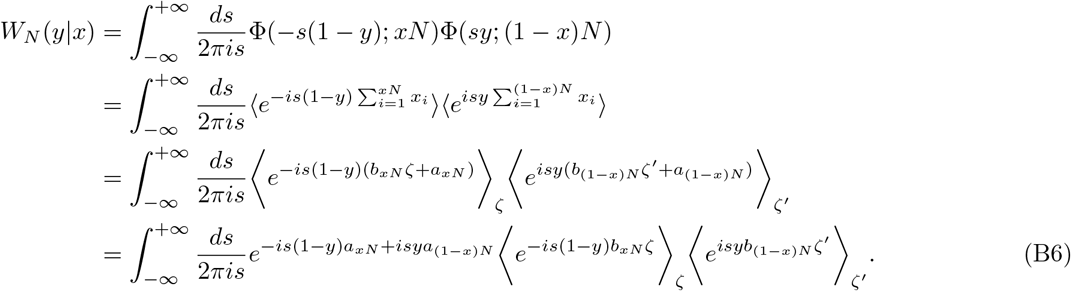

When *N* is large, the quantities in the two brackets in the last line can be approximated by the characteristic functions of *α*-stable distribution, Equation A3, with *s → −s*(1 − *y*)*b_xN_* and *s → syb*_(1−*x*)*N*_, respectively. Thus, when *α* ≠ 1, Equation B6 can be computed as

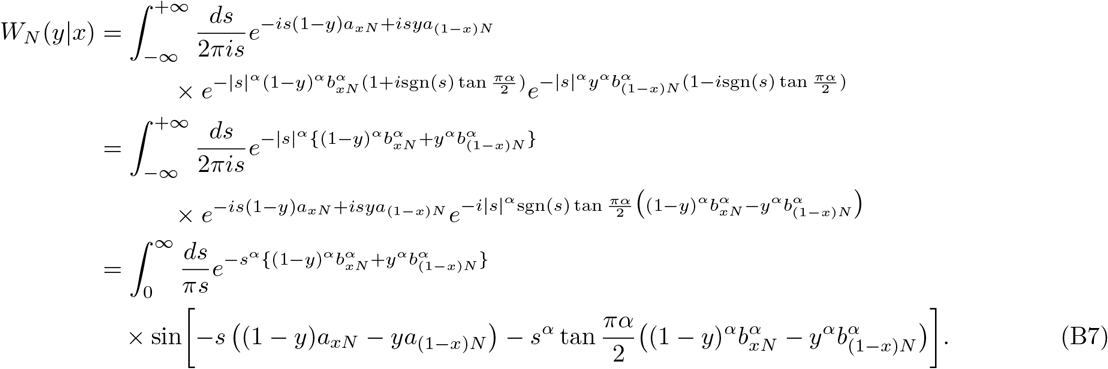

In the following, we evaluate the integral expression of *W_N_*(*y|x*) and compute the transition density *w_N_* (*y|x*) from Equation B3.

**When** *α <* 1

By using Equation A2,

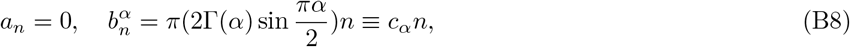

we have

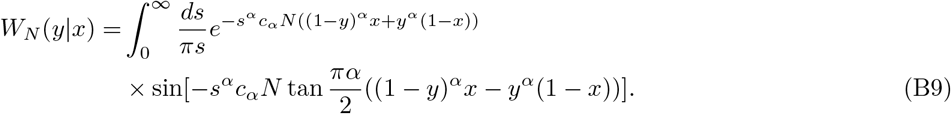

By setting *Nc_α_s^α^* = *σ*, *W_N_* (*y*|*x*) becomes

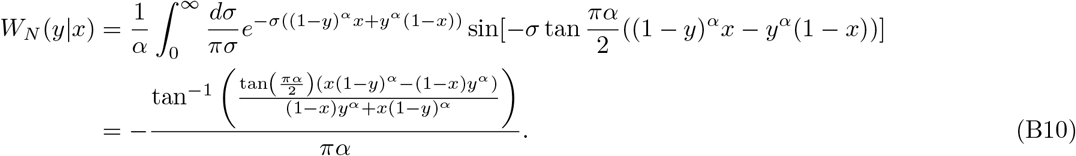

By differentiating it with respect to *y*, we obtain

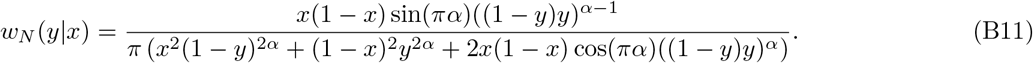

Note that this does not depend on *N*, which is consistent with the fact that the coalescent time is 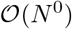 when *α <* 1.

**When** 1 *< α <* 2

By using Equation A2,

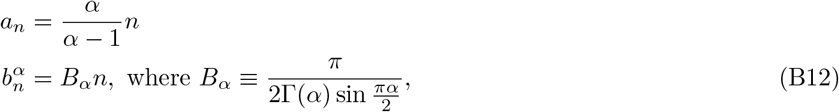

Equation B7 becomes

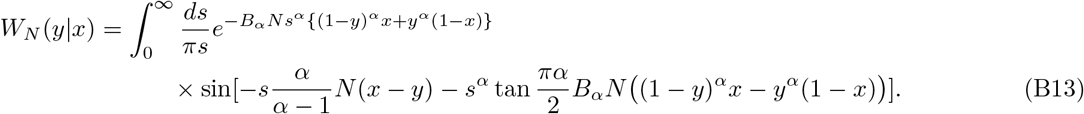

By changing the variable of integration as *σ* = *N* ^1*/α*^*s*, we have

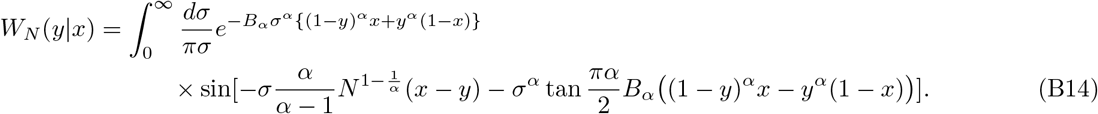

By changing the variable of integration as 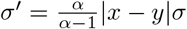 and redefining *σ*′ as *σ*, we have

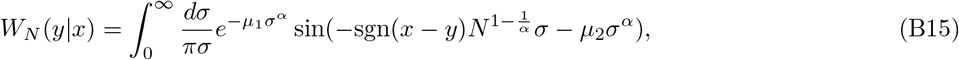

where

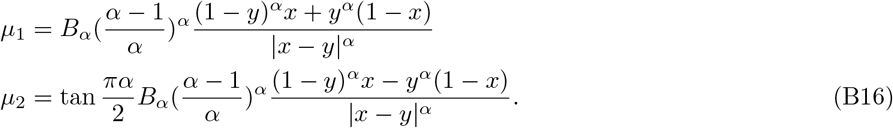

The transition probability *w_N_* (*y*|*x*) is given by

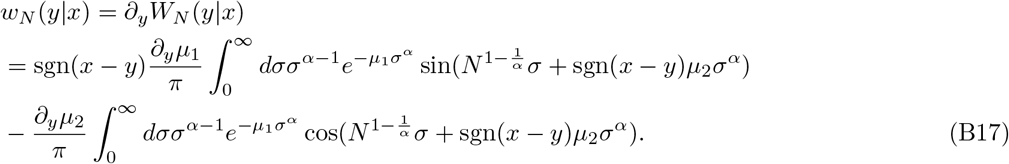

Consider the integral

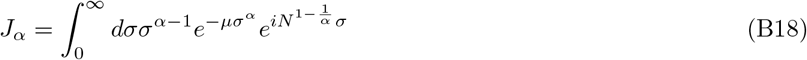

where *μ* = *μ*_1_ − *i* sgn(*x* − *y*)*μ*_2_. Then, the transition probability can be written as

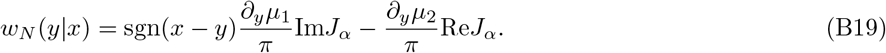

From Watson’s lemma, the integral *J_α_* can be expressed as a series expansion;

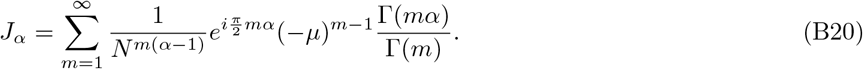

By substituting Equation B20 into Equation B19 and writing *μ* = |*μ*|*e^iθ^*, we obtain

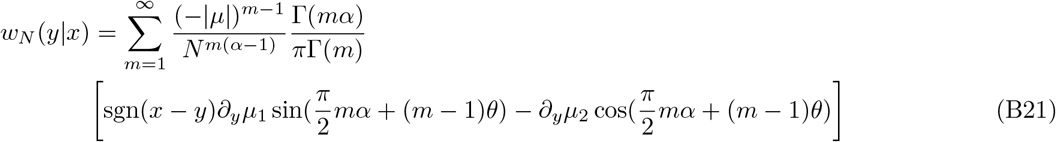

The leading order (*m* = 1) is given by

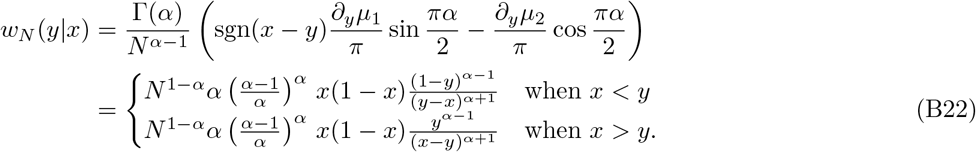

Equation 21 in the main text can be obtained by introducing the continuous time *τ* ≡ *t/*(*C_α_N*^*α*−1^) where 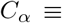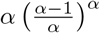. Equation 22 follows from the neutrality 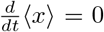. Note that the expansion of Equation B20 is possible only when |*x* − *y*| is finite, i.e., when |*x* − *y*| *> ϵ* where *ϵ* is an *N* -independent positive constant. Although *w_N_* (*y*|*x*) in B22 diverges as |*x* − *y*| → 0, this divergence is not a problem, because the jump term of the asymptotic dynamics in Equation 20 can be obtained from *w_N_* (*y*|*x*) for |*x* − *y*| *> ϵ* (see [37]).

**When** *α* = 2

*a_n_* and *b_n_* are given by

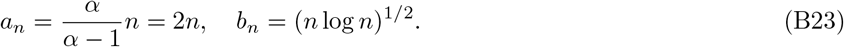

Equation B7 then becomes

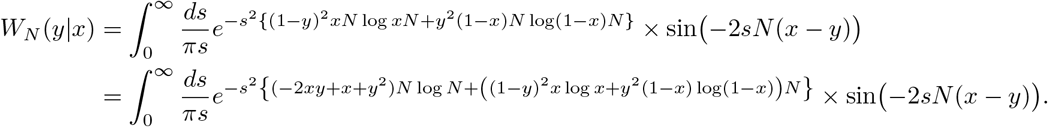

By changing the variable of integration as 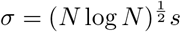,

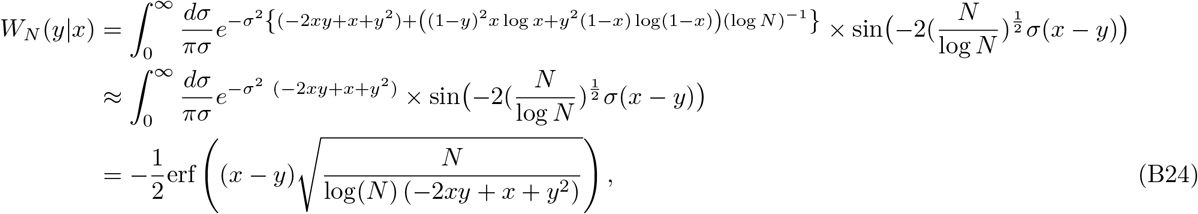

where erf(*x*) is the Gauss error function

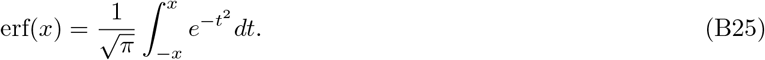

By differentiating *W_N_*(*y*|*x*) with respect to *y*, we have

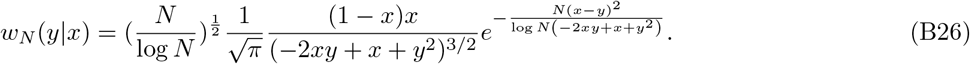

Suppose that *ϵ* is a sufficiently small but finite constant. For |*x* − *y*| *< ϵ*, *w_N_* can be approximated as

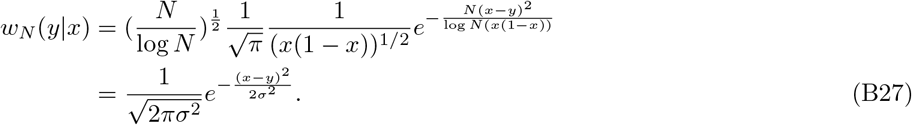

where 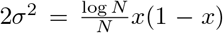. From the symmetry *y − x* → −(*y − x*) of *w_N_* (*y|x*), the advection term is zero. The diffusivity *D* is given by

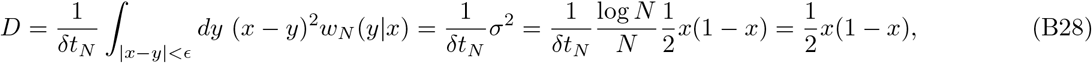

where we have introduced the natural timescale as 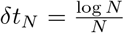 and used the integral approximation

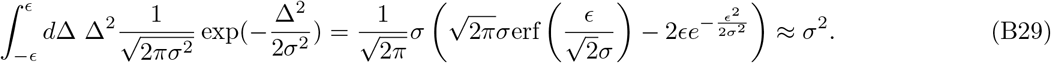

Finally, the jump kernel asymptotically vanishes on the time scale *δt_N_*,

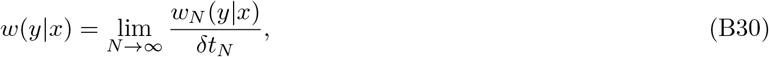

because for fixed *x, y* with |*x − y| > ϵ*, *w_N_* (*y|x*) becomes exponentially small as *N* becomes large.

Thus, in the large-*N* limit, *α* = 2 corresponds to Wright-Fisher diffusion for a population of effective size

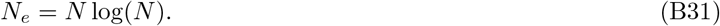

**When** *α >* 2

In this case, since the Pareto distribution 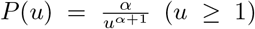 has finite mean 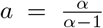 and finite variance 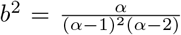, and the large *N* limit of the allele frequency dynamics should be described by the Wright-Fisher diffusion process. To confirm this more generally, we consider a general distribution with finite mean and variance, namely, consider that each individual’s offspring number *u_i_* is sampled from a distribution with mean *a* and variance *b*^2^. Then, from the central limit theorem, the shifted and rescaled variable

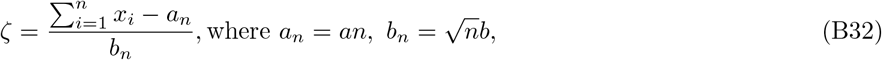

obeys the normal distribution 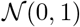. Its characteristic function is given by 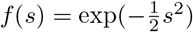. Thus, we have

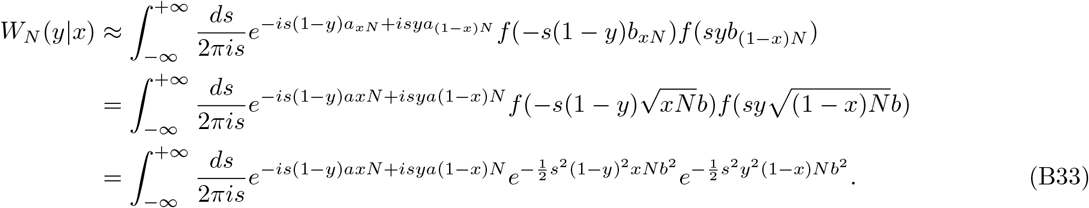

By setting *σ* = *N*^1/2^*s*,

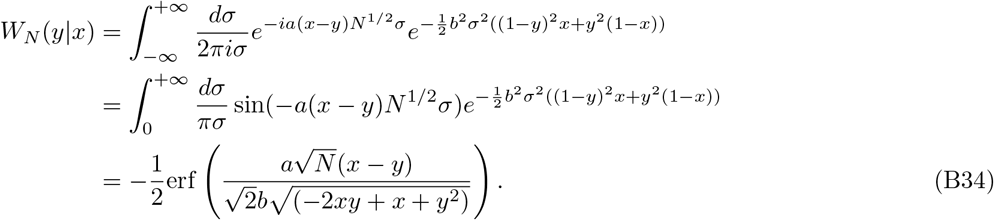

Thus, we obtain

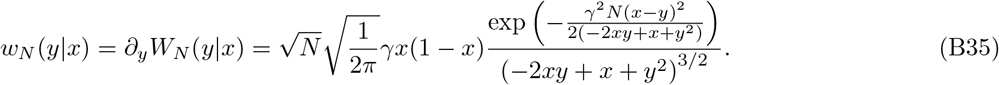

where *γ* ≡ *a/b*. For the Pareto distribution, 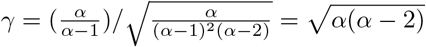.

For |*x* − *y*| *> ϵ*, *w_N_*(*y*|*x*) becomes exponentially small as *N* becomes large, and so the jump term does not exist in the asymptotic dynamics; *w*(*y*|*x*) = 0. For |*x* − *y*| *< ϵ*, we can approximate *w_N_* (*y*|*x*) as

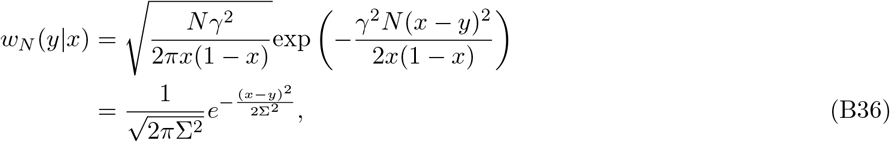

where 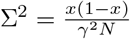. From the symmetry *y* − *x* → −(*y* − *x*) of *w_N_* (*y*|*x*), the advection is zero. Finally, the diffusion is evaluated as

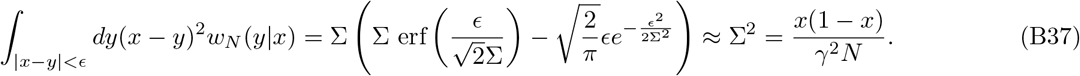

Thus, by re-scaling time as 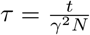, we obtain

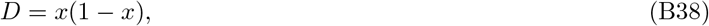

which corresponds to the Wright-Fisher diffusion of a population of effective size *N_e_* = *Nγ*^2^ = *Nα*(*α* − 2). Notice *N_e_* → 0 as *α* → 2, indicating that the concept of the effective population size breaks down when the variance of the offspring distribution diverges.

## Appendix C. The transition density for the differential Chapman-Kolmogorov equation for 1 *< α <* 2

Here we derive the short-time transition density given in Equations 23 and 24 and determine *g*(*ξ*) in the scaling ansatz given in Equation 39.

## 1. The short-time transition density

Before discussing the CK equation in Equation 20, it is instructive to start from the simple diffusion equation,

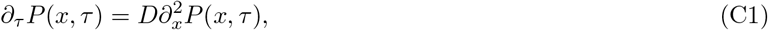

with the initial condition *P* (*x, τ* = 0) = *δ*(*x* − *x*_0_). The solution of this initial value problem is given by

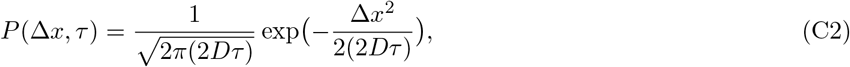

which is usually derived from the Laplace-Fourier transformation. However, this solution can also be obtained by using the central limit theorem: Equation C1 is equivalent to a Brownian motion where jumps *X* → *X* ± *a* occur with rate 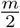, where *a* and *m* are related with *D* via 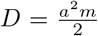. Since *n ≈ mτ* jumps occur in time *τ*, the displacement is approximately given by 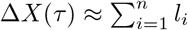 where *l_i_* = ±*a*. Then, from the central limit theorem, Δ*X*(*τ*) is distributed according to the normal distribution with mean *n〈l_i_*〉 = 0 and variance 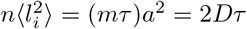, namely, Equation C2. Note that, even if the diffusion constant depends on *x*, the solution in Equation C2 (with *D → D*(*x*_0_)) is valid in short times.

Essentially the same argument can be applied to the CK dynamics, except that the generalized central limit theorem should be employed since the variance of jump sizes is divergent in the case of the CK dynamics. Suppose that the initial density is given by *P* (*x*′, *τ* = 0) = *δ*(*x*′ − *x*) (for notational simplicity, the subscript 0 on *x* is dropped). In the CK dynamics, the frequency change Δ*X*(*τ*) = *X*(*τ*) − *x* is caused by the bias *V* (*x*) in Equation 22 and by stochastic jumps. The rate of a frequency-increasing jump and that of a frequency-decreasing jump are given by

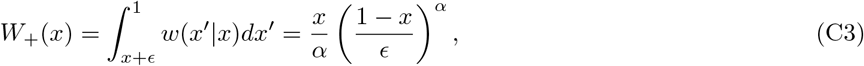

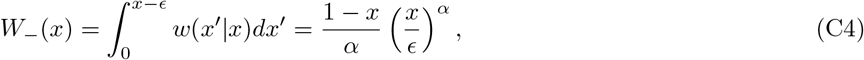

respectively. Therefore, the expected number *n* of jump events in time *τ* is given by

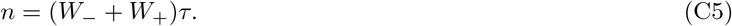

Because randomness in the number of jump events is negligible compared to that in jump sizes, it can be assumed that exactly *n* jumps occur in time *τ*. Then, the displacement Δ*X*(*τ*) = *X*(*τ*) − *x* can be written as

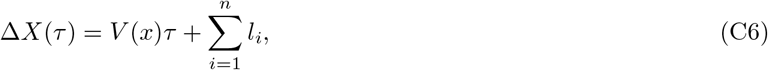

where *l_i_* ∈ [−*x,* −*ϵ*] ∪ [*ϵ,* 1 − *x*] denotes the displacement due to the *i*-th jump. For small *τ*, *w*(*y*|*x*(*τ*′)) ≈ *w*(*y*|*x*) for 0 *< τ′ < τ*, which means that *l*_1_*, …, l_n_* are independent and identically distributed. From Equation 21, each *l_i_* is approximately sampled from the following power-law distribution,

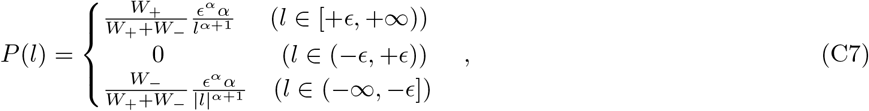

where the factor 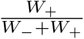 (resp. 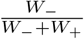) represents the probability that a given jump is frequency-increasing (resp. frequency-decreasing). *P* (*l*) is normalized as 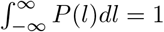. Note that, in Equation C7, the original range [−*x,* −*ϵ*] ∪ [*ϵ,* 1 − *x*] of *l* has been extended to [(−∞ *, −ϵ*] ∪ [*ϵ, ∞*). Under this modification, the variance *x*(*τ*)^2^ is no longer well-defined. However, this modification does not alter short-time properties of typical events, because the presence of the boundaries at *x* = 0, 1 is not important for them.

By noting that *P* (*l*) has a divergent variance and that the number of jumps is 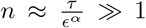 even for small *τ* (as *ϵ* → +0), the generalized central limit theorem states that the sum 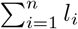 in Equation C6 obeys an *α*-stable distribution. The stable distribution is characterized by 〈*l*〉*, β, γ* given below (see, for example, [42]): The mean 〈*l*〉 is

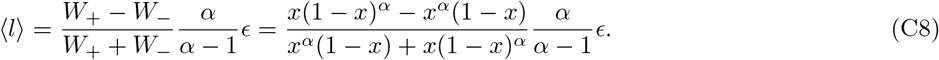

Asymptotically, *P*(*l*) satisfies

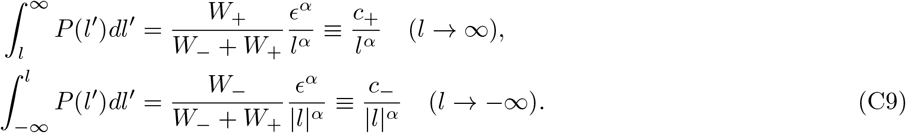

Note *c*_−_ + *c*_+_ = *E^α^*. The parameters *γ* and *β* are determined from *c*_±_;

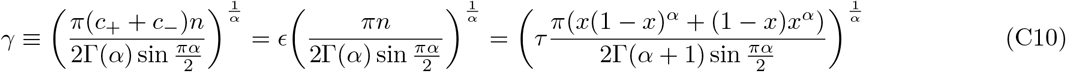

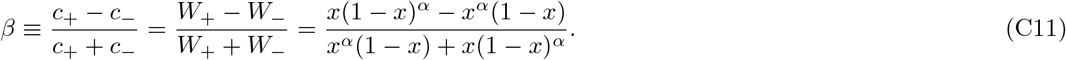

Then, from the generalized central limit theorem, the random variable,

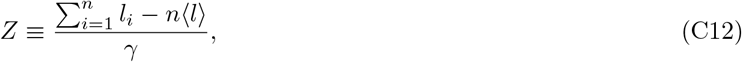

has the following characteristic function,

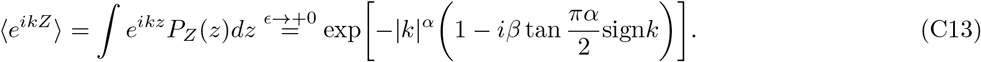

We can determine the characteristic function for Δ*x*, using Equation C13 and the relation

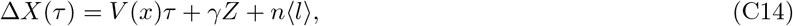

which follows from Equations C6 and C12. While *V* (*x*) and 〈*l*〉 are divergent in the limit *ϵ* → +0, we can show, by using Equation C8 and 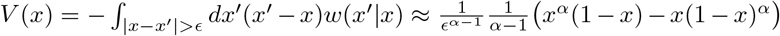, that these divergent terms exactly cancel out each other. Therefore, the displacement is simplified as

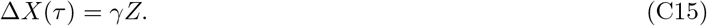

Equations 24 and 23 in the main text are the same as Equations C15 and C13 (with the replacement of *x → x*_0_). By substituting this into Equation C13, we obtain the characteristic function of the allele frequency *X*(*τ*);

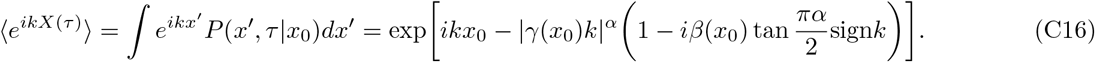

## 2. The scaling ansatz for the long-time transition density in Equation 39

Consider the initial distribution *P* (*x, τ* = 0) = *δ*(*x − x*_0_) with *x*_0_ ≪ 1. After some time, the distribution spreads over the region *x* ≪ 1 with a peak at the extinction boundary *x* = 0. As presented in Equation 39 of the main text, up to a constant prefactor, *P* (*x, τ*) takes the following form

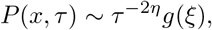

where *η* = (*α −* 1)^−1^ and 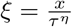. Here, we present an analytic argument to determine *g*(*ξ*).

Equation 20 can be rewritten as

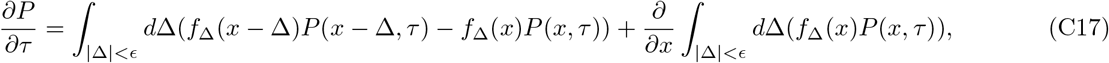

where *f*_Δ_(*x*) ≡ *w*(*x* + Δ|*x*) given by Equation 21. For *x* « 1, *f*_Δ_(*x*) is approximately given by

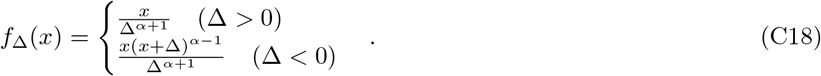

We substitute the ansatz *P* (*x, τ*) ~ *τ*^−2*η*^g(*ξ*) into the above CK equation. The left-hand side of the CK equation becomes

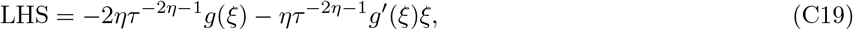

which is proportional to 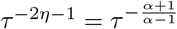. The right-hand side is decomposed into the integrals over Δ *>* 0 and those over Δ *<* 0. We can show that the former is proportional to 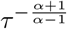, while the latter is proportional to 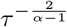; For example, one of the integrals over Δ *>* 0 is

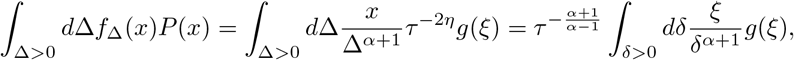

while one of the integrals over Δ *<* 0 is

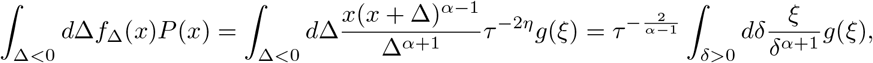

where we have changed the integration variable from Δ to 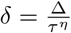. Since the extinction time for the initial frequency *x*_0_ « 1 is much shorter than the coalescent timescale, we can assume *τ* « 1, which implies that the integrals over Δ *>* 0 are negligible compared to those over Δ *>* 0. By evaluating the integrals over Δ *>* 0 using the scaling form of *P* (*x, τ*) and comparing them with Equation C19, we have

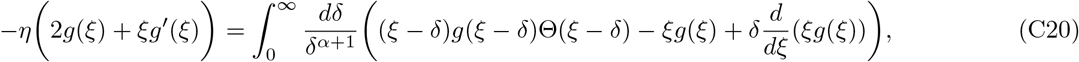

where Θ(·) is the Heaviside step function. Note that the variable of integration has been changed from Δ to 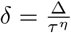, and the upper bound in the integral has been extended into +∞, to make the equation analytically tractable. It is convenient to express Equation C20 in terms of Φ(*ξ*) ≡ *ξg*(*ξ*);

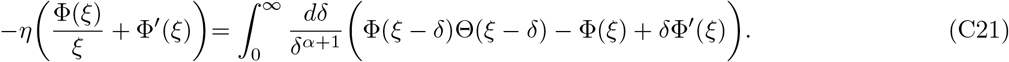

The solution of the integro-differential equation in Equation C21 can be obtained as a series expansion. Assume, for small *ξ*,

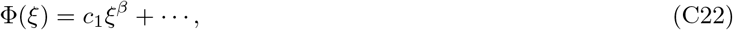

where *c*_1_ is a normalization and the exponent of the leading term is denoted by *β* ∈ (0, 1). Here, *β* < 1 is required since we are considering the situation where *P* (*x, τ*) is monotonically decreasing in *x*, while *β* > 0 is required to make *P* (*x, τ*) normalizable. By substituting Equation C22 into Equation C21, we have

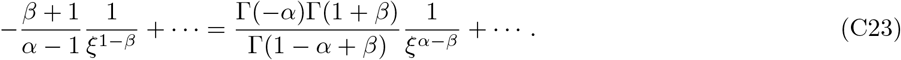

Since 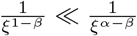 for *ξ* ≪ 1, in order for the two sides to be balanced, the coefficient 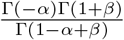 needs to be zero, which is possible only when Γ(1 − *α* + *β*) diverges. Since 1 < *α* < 2 and 0 < *β* < 1, we can conclude *β* = *α* − 1. Therefore, the leading term of *g*(*ξ*) is given by

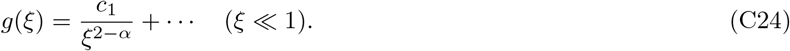

More generally, by starting from the ansatz,

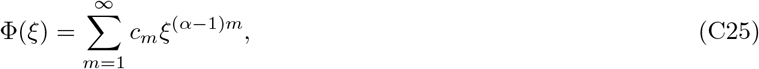

the coefficients *c*_2_, *c*_3_, · · · can be determined iteratively:

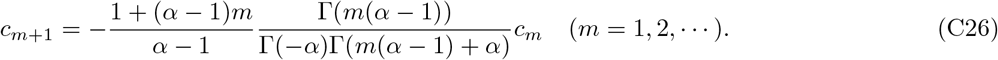

By using this iteratively, we can express Φ(*ξ*) as

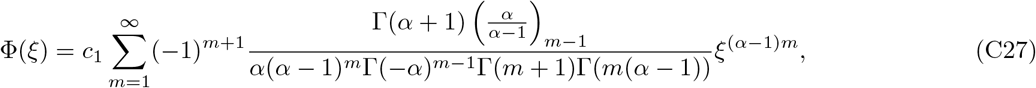

where 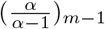 (is the Pochhammer symbol, (*q*)_*n*_ = Γ(*q* + *n*)/Γ(*q*). The analytic expression of *g*(*ξ*) can be obtained from this using 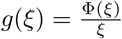.

On the other hand, for *ξ* ≫ 1, we expect that *g*(*ξ*) decreases in the same way as the offspring distribution does;

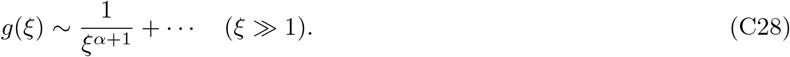

Therefore, we expect there is a crossover point *ξ*_c_ such that 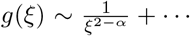 for *ξ* ≪ *ξ*_c_ and 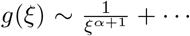 for *ξ* ≫ *ξ*_c_. The scaling form for *ξ* ≫ *ξ*_c_ can indeed be confirmed by considering the following ansatz for Φ(*ξ*),

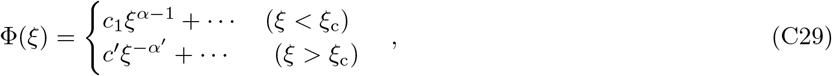

where *c*′ is a normalization and *α*′ is an exponent to be determined. Substituting this ansatz into Equation C21, we can show *α*′ = *α*, leading to 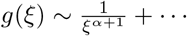 for *ξ > ξ*_c_.

Finally, we remark that, while Equation C27 is derived assuming *ξ* ≪ 1, the series converges for any *ξ* > 0. This indicates that the scaling form 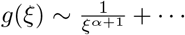 for large *ξ* should directly follow from a resummation of the infinite series in Equation C27. In fact, numerical evaluation of a finite truncation of the series indicates the crossover behavior Equation C29 (see Figure 15).

**Figure 15.**
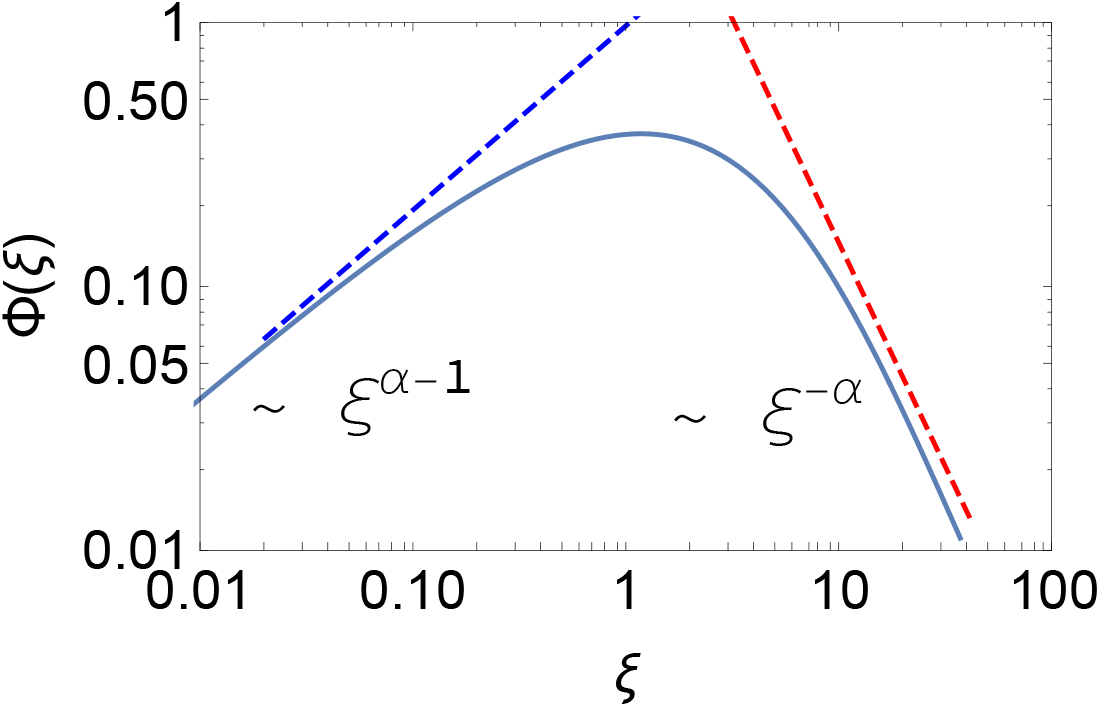
The infinite series in Equation C27 is evaluated numerically by truncating at *m* = 150 and using the van Wijngaarden transformation (solid line). *α* = 1.7 is used. The dashed blue and red lines represent the asymptotic behaviors given in Equation C29.

## Appendix D. From Lambda-Fleming-Viot Generator to differential Chapman-Kolmogorov equation

In Appendix B, the jump density *w*(*y|x*) is derived from the generalized Wright-Fisher sampling, Equation 13 in the main text. Here, we present another more formal derivation of the jump density *w*(*y|x*) for 1 < *α* < 2. See [32] for the case *α* = 1.

## 1. Jump density for general Λ measure

The backward generator of the Λ coalescent process for the biallelic model (see e.g. [43, 44]) is given by

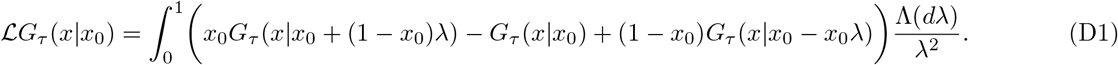

This can be rewritten as a sum of two terms:

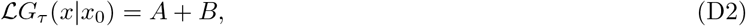

where

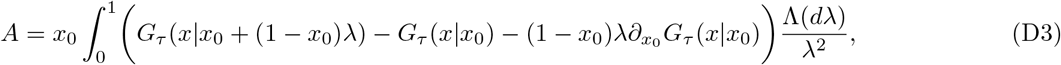

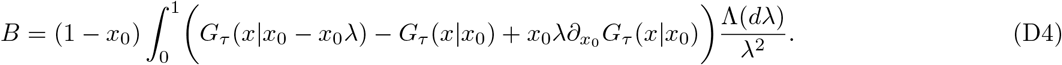

We introduce the integration variable *x*′ ≡ *x*_0_ + (1 − *x*_0_)*λ* for *A* and *x*′ ≡ *x*_0_ − *x*_0_*λ* for *B* respectively. By writing

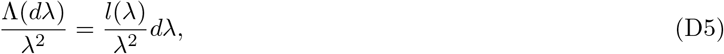

*A* and *B* become

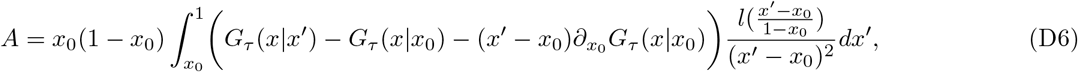

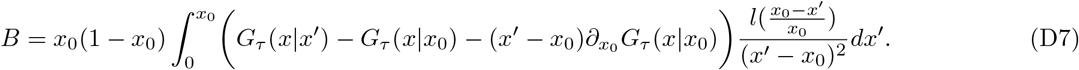

Defining the jump kernel *w*(*x*|*x*_0_) as

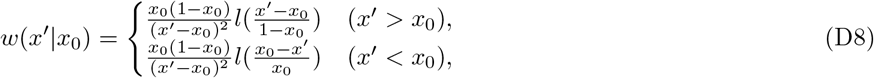

we can formally rewrite the generator as

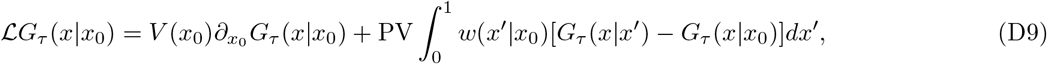

where

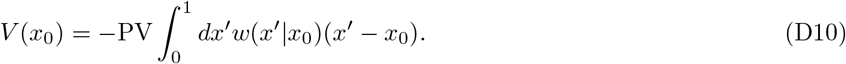

## 2. When the measure is the Beta distribution Beta(*α,* 2 − *α*)

We take the Beta(*α,* 2 − *α*) distribution as the Λ measure, which corresponds to the descendant distribution considered in this study, ~ 1*/u*^1+*α*^:

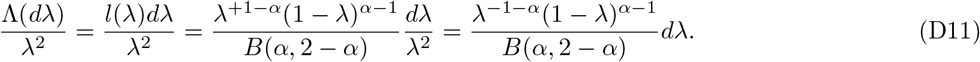

With this measure, *A* and *B* become

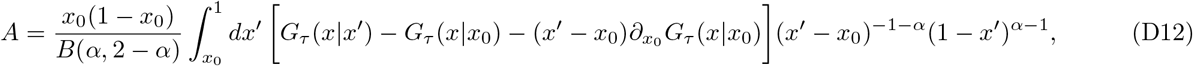

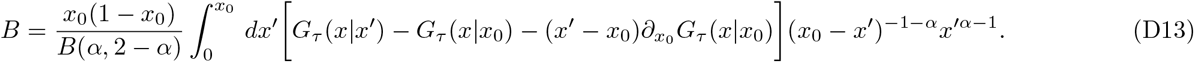

Note that the integrals *A* and *B* are convergent for *α* ∈ (0, 2), because, near *x*′ ~ *x*_0_, the terms inside [···] are 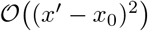 and so the integrands are 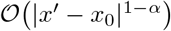. The jump kernel is given by

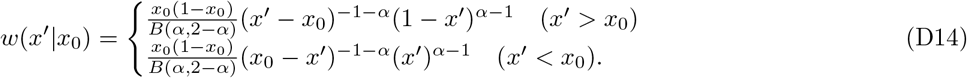

When 1 < *α* < 2, this density agrees with Equation 21 of the main text (up to a proportionality constant). The advection is given by

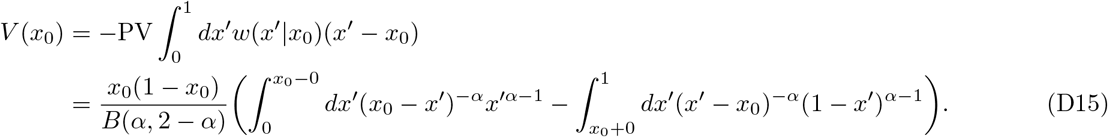

Note that, when *α* > 1, the limit 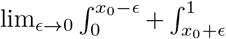 in Equation D15 does not exist, although this divergence is rather formal since there exists a natural cutoff 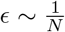 for a finite-size population.

## Appendix E. Analytic results in the marginal case *α* = 1

Although the main target of our present study is the case of 1 < *α* < 2, we here provide analytical results for *α* = 1, which have not been derived before.

## 1. Site frequency spectrum in the presence of genuine selection

The transition density for *α* = 1 in the presence of natural selection is derived in [32] (see [27] for neutral case). In

*x* space, it is given by

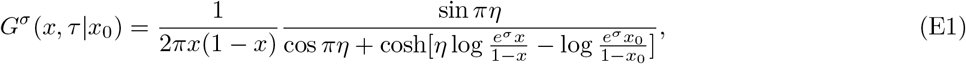

where *η* ≡ *e*^−*τ*^ and *σ* is the selective advantage (there is an erratum in Equation 38 in [32]).

For the purpose of computing the site frequency spectrum (or, equivalently, the mean sojourn time), we set *x*_0_ = 1*/N*. Since we are considering the large *N* limit, the denominator of Equation E1 can be rewritten as

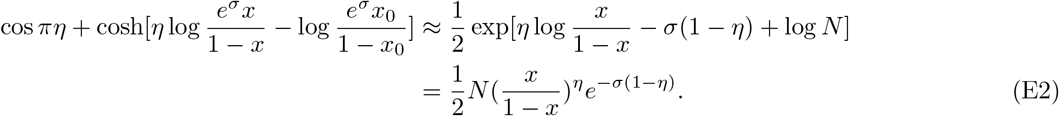

Thus, the transition density for 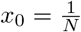 can be written as

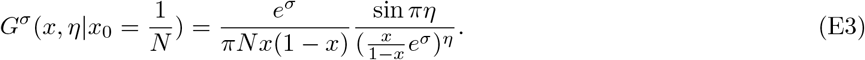

Near the boundaries, this can be approximated as

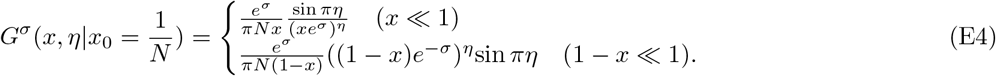

The site frequency spectrum is given by *f*_SFS_(*x*) = *Nμ × t*(*x*), where *μ* is the mutation rate per generation, and *t*(*x*) is the mean sojourn time density, which is given by

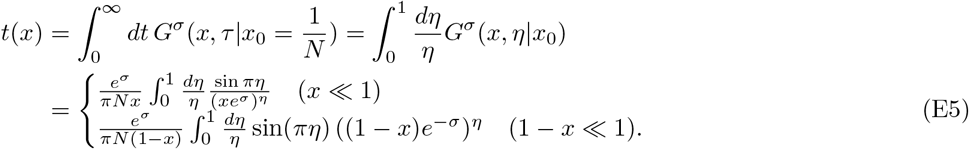

Next, we compute the integrals in Equatiion E5, asymptotically close to the absorbing boundaries (see Equation E14 for the final results). To evaluate Equation E5 for *x* ≪ 1, we first consider the integral,

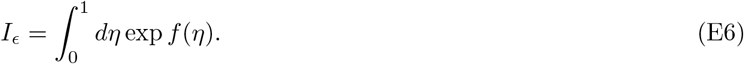

When *f* (*η*) has a sharp peak at *η* = *η*^*^, we approximate this integral a

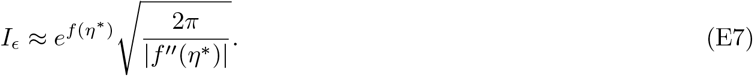

In our case,

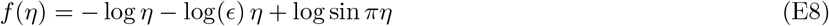

where *E* = *xe^σ^*. *f* (*η*) takes the maximum value at 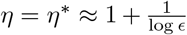^1^. At *η* = *η*^*^, 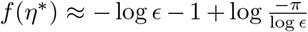, and 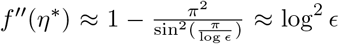. The saddle-point evaluation in Equation E7 is precise when *ϵ* ≪ 1. By using these expressions, *I_ϵ_* can be evaluated as

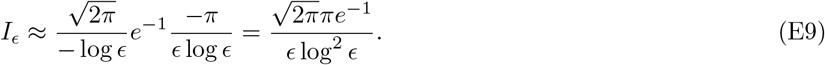

By setting *E* = *xe^σ^*, we find

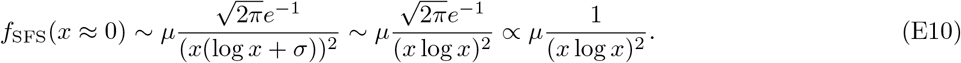

Next, to evaluate Equation E5 for the high-frequency end, we consider the following integral

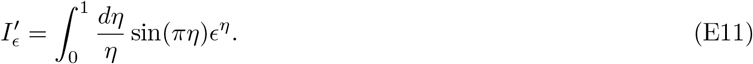

When *ϵ* ≪ 1, the integrand takes the maximum value at the boundary *η* = 0. Thus,

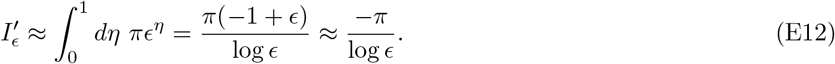

By setting *ϵ* = (1 − *x*)*e*^−*σ*^, we find

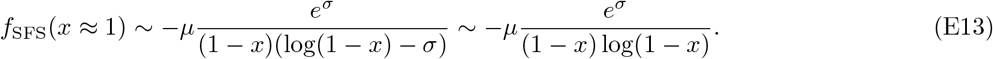

In summary, the SFS in Equation E5 is given by

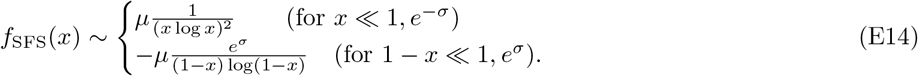

Note that the dependence on *σ* disappears when *x* ≪ 1. Figure 6 shows the plots of the SFS.

For comparison, we write the site frequency spectrum for the Wright-Fisher model (*α* ≥ 2) (see, for example, [3, 51]);

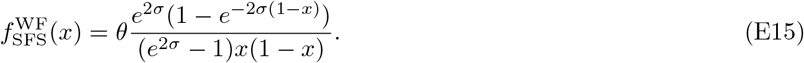

The asymptotic forms near the boundaries are given by

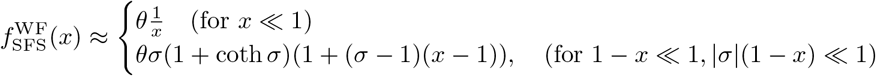

where we have expanded the SFS around *x* = 1 up to the sub-leading order. For a sufficiently strong selection (*σ* > 1), the SFS increases with *x* at the high-frequency end. However, unlike the case of *α* < 2, the increase is not strong and the SFS approaches the constant *σ*(1 + coth*σ*) as *x* → 1.

## 2. Dynamics of the median of allele frequencies

When *α* = 1, we can derive a simple differential equation that described the median of trajectories. In the logit space, the transition density is given by

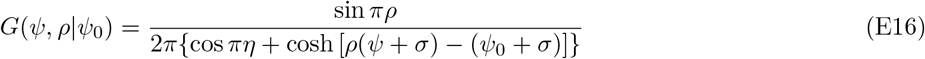

where *ρ* = *e*^−*τ*^. The median Ψ^med^ (at a given time point *ρ*) is characterized by

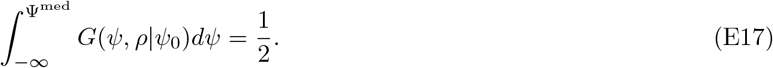

From the symmetry of cosh, the median is given by the peak of the transition density;

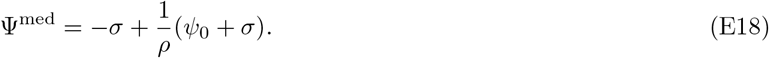

By differentiating Equation E18 with respect to *ρ* and eliminating *Ψ*_0_, we obtain

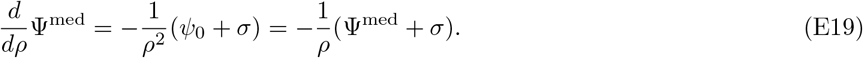

Noting that 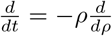, we find

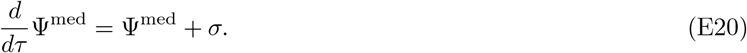

Since the median is invariant under a coordinate transformation, the median *X*^med^ in the *x* space is simply related with Ψ^med^ via the logit transformation, 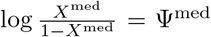. By differentiating this with respect to time and using Equation E20, we obtain

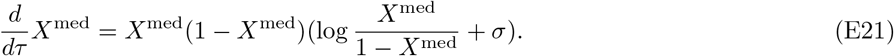

## 3. Allele frequency dynamics conditioned on fixation

By using Bayes’ theorem, the probability distribution of the allele frequency conditioned on fixation can be written as

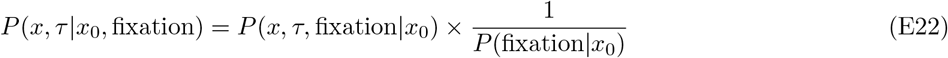

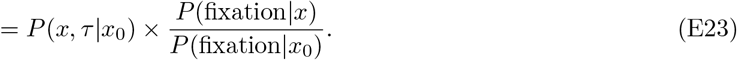

The fixation probability for the initial frequency *x*_0_ is given by (see [32])

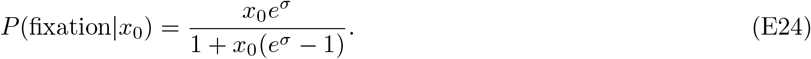

In particular, the fixation probability of a single mutant is given by

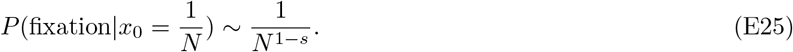

By using Equation E24, the conditioned probability in Equation E23 is computed as

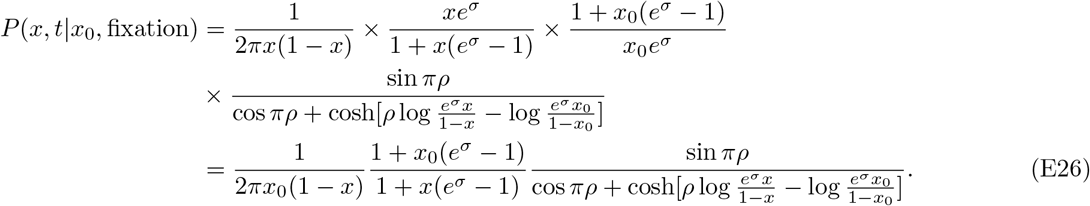

## Appendix F. Site frequency spectra in presence of selection

Here, we argue the effect of the genuine selection on the SFS by using the effective bias when 1 < *α* < 2. As discussed in the main text, there is a crossover point *x_c_*, shown in Equation 41, below which the selection is negligible compared to the effective bias (see Figure 13). Thus, we can expect that the SFS becomes independent of the selective advantage *σ* for a sufficiently small frequency *x*. Similarly, for the high-frequency end 1 − *x* ≪ 1, the selection is negligible compared with the effective bias. Therefore, we expect that 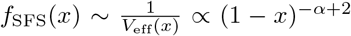 even in the presence of natural selection. In particular, the exponent is independent of *σ*. Figure 17 shows the numerical results when *α* = 1.5. As *x* approaches 0, the SFS becomes independent of the selective advantage *σ*. For frequent variants 1 − *x* ≪ 1, the SFS can be fitted well by (1 − *x*)^−*α*+2^, while the magnitude of the SFS increases with *σ*. A similar result can be obtained analytically when *α* = 1 (see Appendix E).

**Figure 16.**
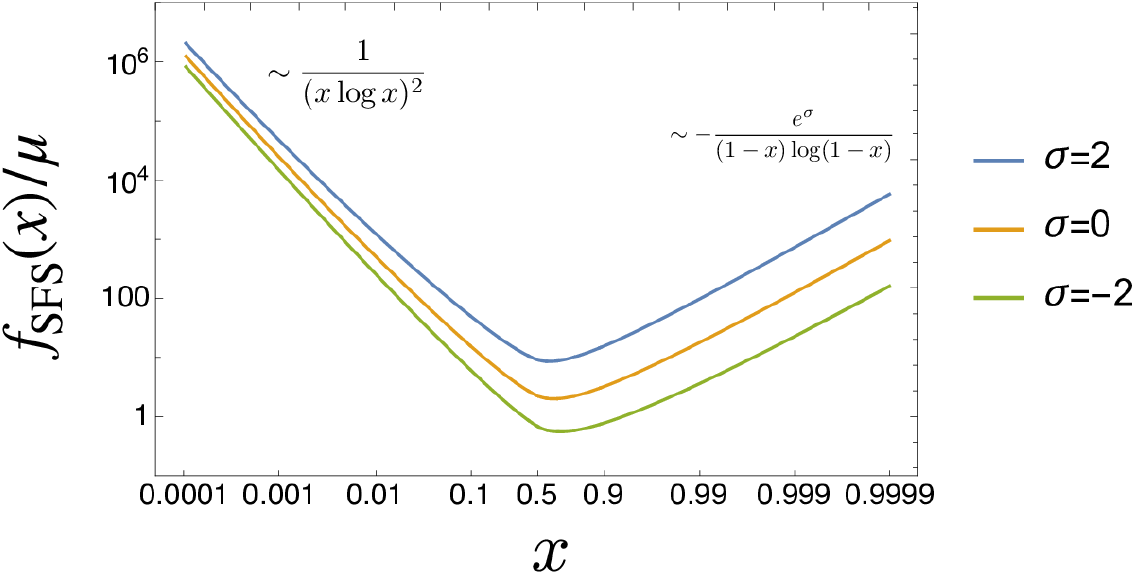
The SFS *f*SFS(*x*)*/μ* when *α* = 1 for the selective advantage *σ* = −2, 0, 2. *f*SFS(*x*) is obtained by numerically evaluating the exact expression of *t*(*x*) in the first line of Equation E5. As *x* ⟶ 0, *f* (*x*) becomes independent of *σ*. Near *x* = 1, while the magnitude of *f* (*x*) depends on *σ*, the scaling behavior (slope in the log-log plot) does not. See Equation E14.

**Figure 17.**
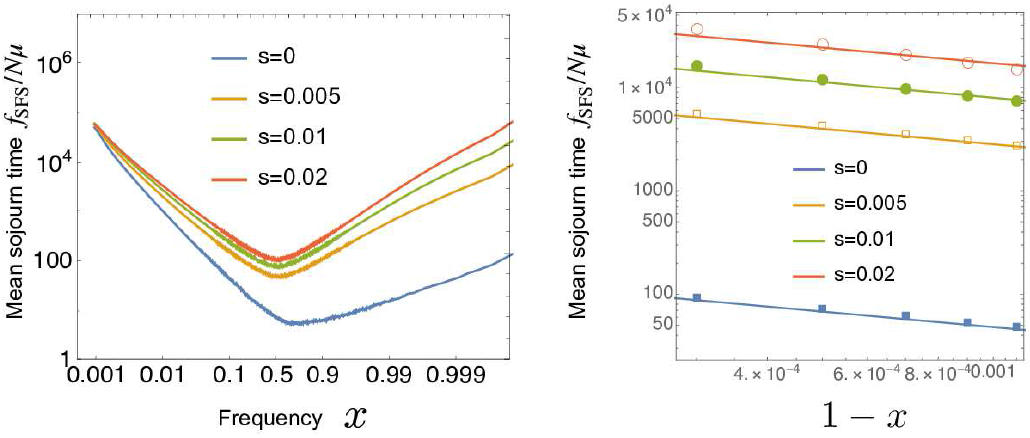
Left: The SFS under positive selection *s* = 0, 0.005, 0.01, 0.02. *α* = 1.5 and *N* = 10^6^. Right: The SFS near *x* = 1. The straight lines are drawn assuming *SFS*(*x*) ∝ 1/(1 − *x*)^2−*α*^. The slope is almost independent of *s*.

## Appendix G. Derivation of the rate of adaptation in Equation 48 of the main text

Here, we conjecture the rate of adaptation for an asexual population with a broad offspring distribution (1 < *α* < 2) in the clonal-interference regime, using a self-consistency condition argument described in [48].

We assume that mutations have a fixed effect *s* much larger than the mutation rate *μ*_B_ at which they arise. First, we consider the dynamics of the fittest sub-population that becomes established at the nose of the fitness wave. We can estimate the size of the sub-population when established from the establishment probability of a single fittest mutant;

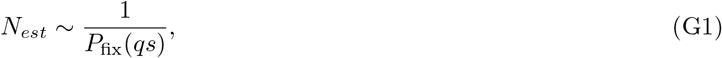

where *qs* (*q* ∈ ℕ) is the fitness lead of the sub-population compared with the mean of the whole population, and the fixation probability is given by Equation 42, 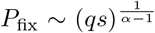. In the time this sub-population is seeded and becomes established, the mean fitness should increase by *s*. This implies that, after its establishment, this sub-population will initially grow exponentially at rate (*q* − 1)*s*. The growth rate will slow down to 0 when it fixes. Therefore, the time from establishment to fixation can be estimated as

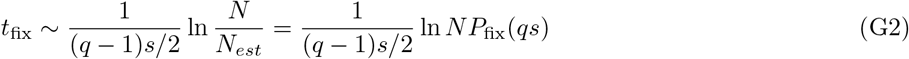

where (*q* − 1)*s/*2 is its average growth rate between the establishment and fixation. Thus, the rate of adaptation is given by

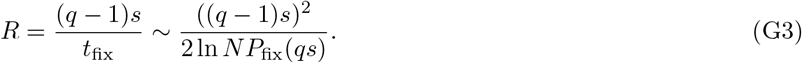

Second, we focus on successive events of establishments at the edge of the fitness wave. We define *t_est_* as the mean time interval between two successive establishments. An established sub-population grows like 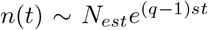, from which the next event of establishment is produced with rate *n*(*t*)*μ*_B_*P*_fix_(*qs*). Therefore, *t_est_* can be estimated from

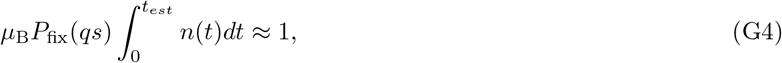

which leads to 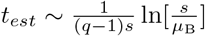. Since the nose of the fitness wave advances at a speed 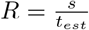, we have

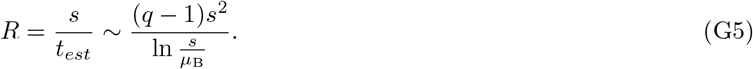

By comparing Equations G3 and G5, we obtain

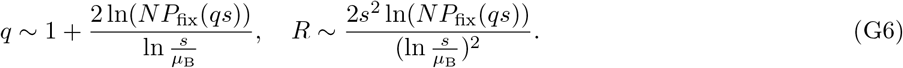

By substituting 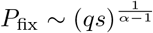 into Equation G6, we obtain

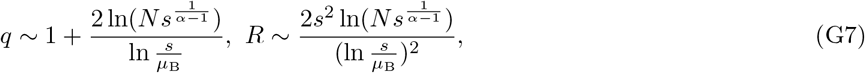

where we used 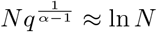. In the limit *α* → 2, the above results reproduce those in [48].

The case of *α* = 1 can be discussed in a similar way. Suppose that the population is monoclonal. The fixation probability of a mutant is given by *P*_fix_ ~ *N*^−1+*s*^ (see Equation E25), which implies that the establishment size is roughly given by *N_est_ ~ N* ^1−*s*^. While the timescale of establishment of a mutant is given by (*μ*_B_*NP*_fix_)^−1^ = (*μ*_B_*N^s^*)^−1^, the timescale of fixation is given by 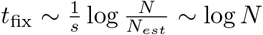. Thus, the successive selection sweeps occur if (*μ*_B_*N^s^*)^−1^ ≫ log *N*, or equivalently,

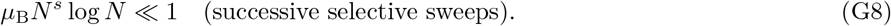

By substituting *P*_fix_ ~ *N* ^−1+*s*^ into Equation G6, the rate of adaptation in the clonal-interference regime is given by

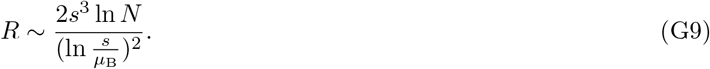

In the successive-sweeps regime, the adaptation rate is given by

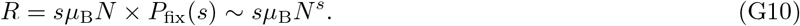

Note that clonal interference becomes unlikely to occur as the offspring distribution becomes broader. For example, when *α* = 1, the population size needs to be *N* ≫ 10^41^ for *μ*_B_ = 10^−4^*, s* = 0.05 to satisfy *μ*_B_*N^s^* log *N* ≫ 1.

Figure 18 shows the numerical results of the adaptation rate *R* versus the selection coefficient *s*. The parameters used in the simulation are in the regime of clonal interference. When 1 < *α*, *R* is approximately proportional to *s*^2^, while, when *α* = 1, *R* is approximately proportional to *s*^3^, which are consistent with Equations G6 and G9. However, when *α* = 1, the quantitative agreement between the numerical result and the theoretical prediction is not good, and a further investigation is needed to validate Equation G9.

**Figure 18.**
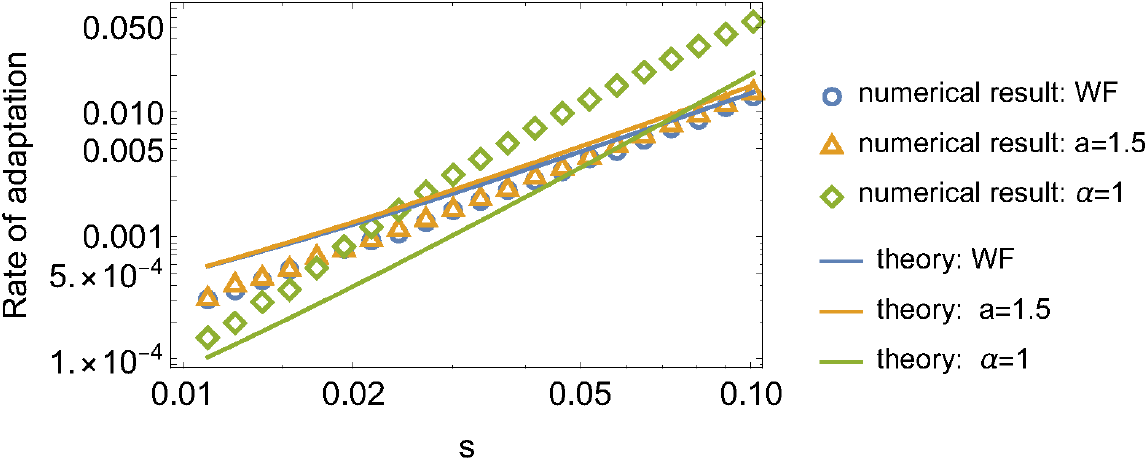
The open markers show the numerical results of *R* as a function of *s*, while the curves show the theoretical predictions, based on the heuristic argument. The m rate of beneficial mutations is *μ* = 10^−4^. The population size is *N* = 10^100^ for *α* = 1,*N* = 10^10^ for *α* = 1.5, and *N* = 10^8^ for the Wright-Fisher model.

## Appendix H. Stationary distributions of traveling wave model in the presence of natural selection

In Figure 14 of the main text, the mutant allele is assumed be neutral. Here, we provide the results in the case where mutants have a fitness advantage *σ* (Figure 19). As in the main text, symmetrically reversible mutations are assumed.

**Figure 19.**
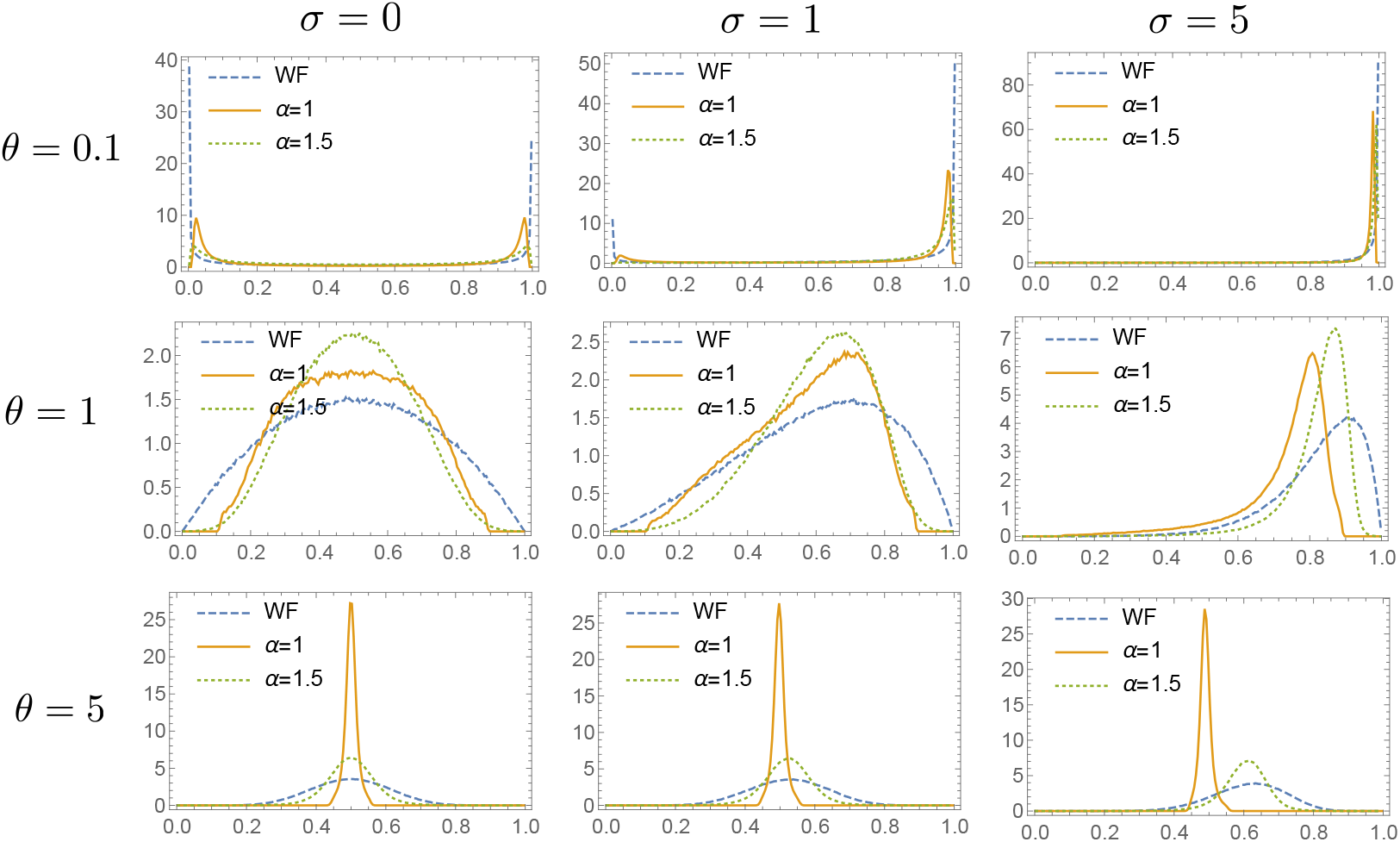
The stationary distributions of the mutant frequency for *θ* = 0.1, 1, 5. *σ* = 0, 1, 5. *σ* is the selection coefficient in the time-continuous description, *σ* = *sT_c_*.

## Appendix I. Numerical simulations

Simulations are implemented in C++ with the GNU scientific library’s random number generators. Results obtained from the simulations are analyzed by Mathematica. The codes are freely available upon request.

## 1. Numerical synthesis of Pareto random variables and *α*-stable distribution

In order to generate the mutant frequency of the gamete pool, we need to compute the sums of random Pareto variables,

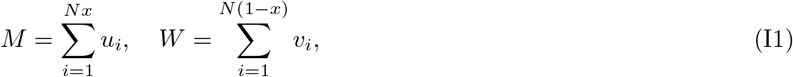

where *u_i_, v_i_* are drawn from the Pareto distribution *P_U_*(*u*) = *α/u^α^*^+1^ (*u* ≥ 1). One simple way to synthesize *u_i_, v_i_* is to sample a number *r* from the uniform distribution on (0, 1) and compute 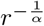.

To generate the sums *M, N* efficiently for large *N* (e.g. *N* ~ 10^6^), we can use the generalized central limit theorem when *xN* and (1 − *x*)*N* are large. In simulations, when *xN* < 100, *M* is generated directly by synthesizing *xN* random variables {*u_i_*}, while, when *x*L*N* ≥ 100, *M* is generated by sampling a random number *ζ* from the *α*-stable distribution and then determining *M* = ∑_*i*_*u_i_* from Equation A1. *W* is generated in a similar way.

After generating *M* and *W*, the population is updated by the binomial sampling with the success probability 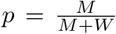 (although this sampling process can be omitted when *α* ≥ 2 since the fluctuations associated with the binomial sampling is negligible compared to the fluctuations associated with *M* and *N*). Natural selection and mutations are implemented by modifying the success probability 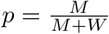 as

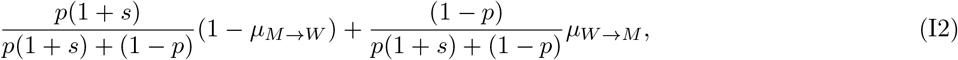

where *μ_W_*_→*M*_ is the mutation rate from the wild-type to the mutant allele, and *μ_M_*_→*W*_ is the mutation rate in the reverse direction.

## 2. Site frequency spectrum

Since the SFS is proportional to the mean sojourn time, the SFS can be computed numerically by generating trajectories staring with 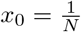 until fixation or extinction and measuring how many times a trajectory visits a given frequency interval on average.

## 3. Numerical simulation of the model of range expansion in the main text

We first review the numerical implementation of the range expansion model with two neutral alleles without mutations [21]. The per capita growth rate *r*(*n*) with an Allee effect is given by

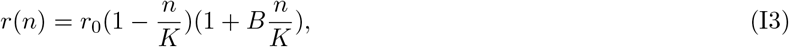

where *n* = *n*_1_ + *n*_2_ is the sum of the two population densities, and *B* is the strength of cooperativity. In each deme, there are three types; allele 1, allele 2, and “empty”. At each time step, the configuration of deme *x* is updated by the trinomial sampling process with

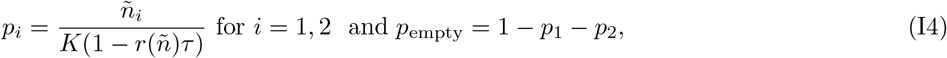

where 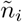 is the population density after migration,

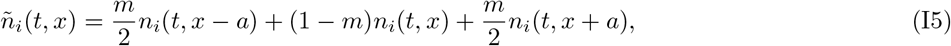

and 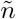 in the denominator of Equation I4 is the sum of these densities, 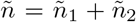, and *a* denotes the width of a deme. The expectation value of the total density *n* after one time step is given by

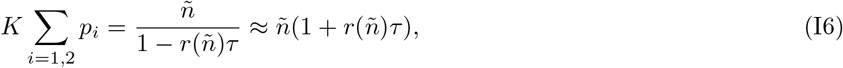

which explains the denominator of Equation I4. In the simulation, *a* = 1 and *τ* = 1 are used.

As in the standard Wright-Fisher model, a mutation process can be introduced by using the success probabilities 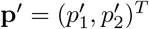 given by

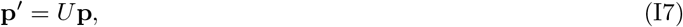

where **p**= (*p*_1_*, p*_2_)^*T*^ and *U* is a matrix representing mutational transitions. In the case of symmetrical mutations in the main text, *U* is given by

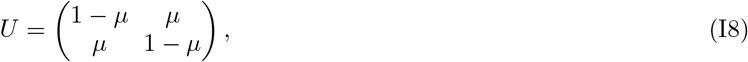

This model serves as a microscopic description of our (non-spatial) macroscopic model of the population with a broad offspring distribution 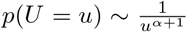. We can argue the relation between the parameters in the two models by comparing the coalescent timescales. As established in [21], for a semi-pushed wave (2 < B < 4), the coalescent timescale is given by

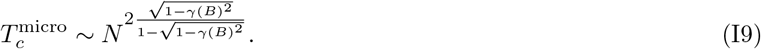

where 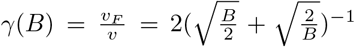 is the ratio of the Fisher velocity 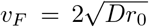 to the wave velocity 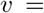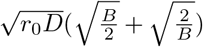. On the other hand, the coalescent timescale 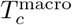 in the macroscopic description for 1 < α < 2 is proportional to *N^α^*^−1^ (see Equation 15). By comparing the exponents, a semi-pushed wave with *B* corresponds to the macroscopic model with ^3^

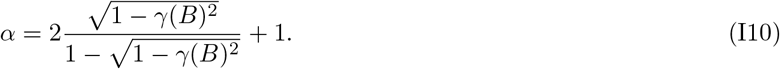

For example, *B* = 3 corresponds to *α* = 1.5. In addition, the mutation rate *μ*_micro_ per generation in the microscopic model and the mutation rate *μ*_macro_ per generation in the macroscopic model should be related by 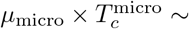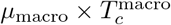.

In the three panels (Left. Center, Right) in Figure 14B of the main text, The following parameters are used.

- Left: *B* = 1*, μ* = (5 × 10^−4^, 5 × 10^−5^)*, K* = 28000 for the microscopic model model, and *α* = 1*, θ* = (1.5, 0.15) for the macroscopic model.
- Center: *B* = 3*, μ* = (2 × 10^−4^, 2 × 10^−5^)*, K* = 35000 for the microscopic model, and *α* = 1.5*, θ* = (1.6, 0.16) for the macroscopic model.
- Right: *B* = 8*, μ* = (1 × 10^−5^, 1 × 10^−6^)*, K* = 57000 for the microscopic model, and the Wright-Fisher model, *θ* = (2.4, 0.24) for the macroscopic model.

In all of the three cases, the growth rate 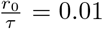 and the migration probability 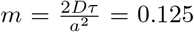 are used in the microscopic model, and the population size *N* = 10^5^ is used in the macroscopic model. Note that, to compare the microscopic model with the macroscopic model, the value of the carrying capacity *K* for each case is chosen such that the size of the front population 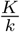, where *k* is the spatial decay rate of the population density^4^, approximately agrees with the population size *N* = 10^5^ in the macroscopic model.

## Appendix J. Areas swept by trajectories

## 1. A scaling argument on area distributions

Consider frequency trajectories that depart from a single mutant 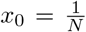 and are eventually absorbed either at *x* = 0 or at *x* = 1. For each of such trajectories, we can define the area in *x* − *τ* -space swept by the trajectory (see Figure 20),

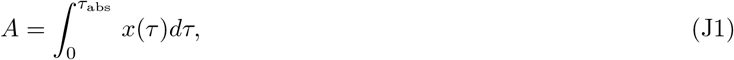

where *τ*_abs_ is the absorption time of the trajectory. While this quantity is defined for a population without spatial structure, we expect that it has a natural interpretation in a model of range expansion as a spatial integration over the mutant frequency (i.e., the abundance of the mutant type), since *τ* in Equation J1 is related with the spatial position of the traveling wave in the comoving frame.

**Figure 20.**
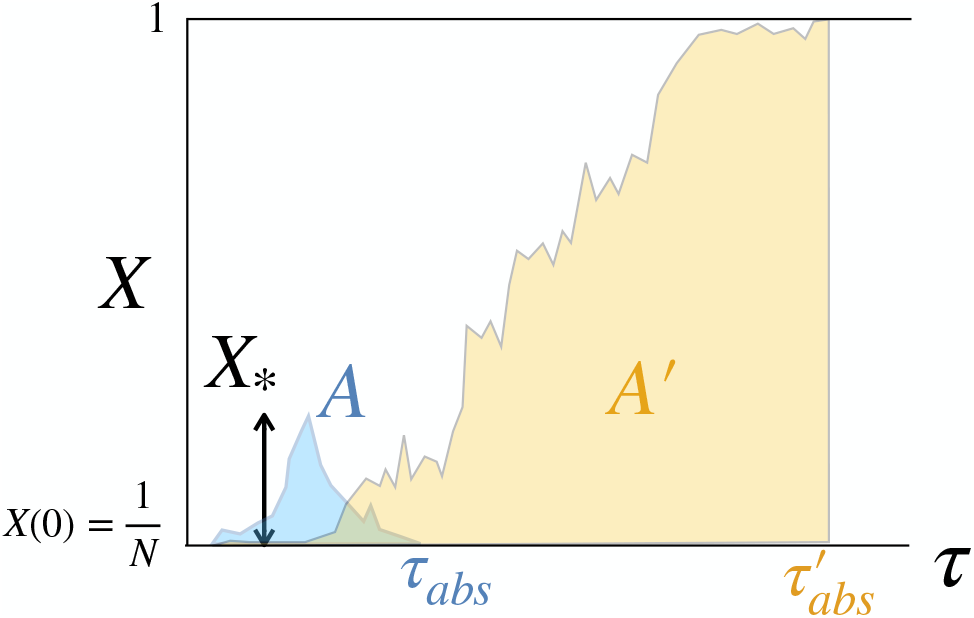
An area *A* swept by a trajectory that eventually goes extinct and an area *A*′ swept by a trajectories that eventually gets fixed are illustrated. *τ*_abs_ and 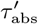 are the extinction time and the fixation time, respectively.

Here, we examine how the area *A* defined in Equation J1 depends on the exponent *α* of the offspring distribution. The left panel of Figure 21 shows the numerical results of the area distribution *p*(*A*) for *α* = 1, 1.5, and the Wright-Fisher model (corresponding to *α* ≥ 2). In a wide range of *A*, areas are distributed according to 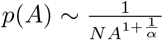.

**Figure 21.**
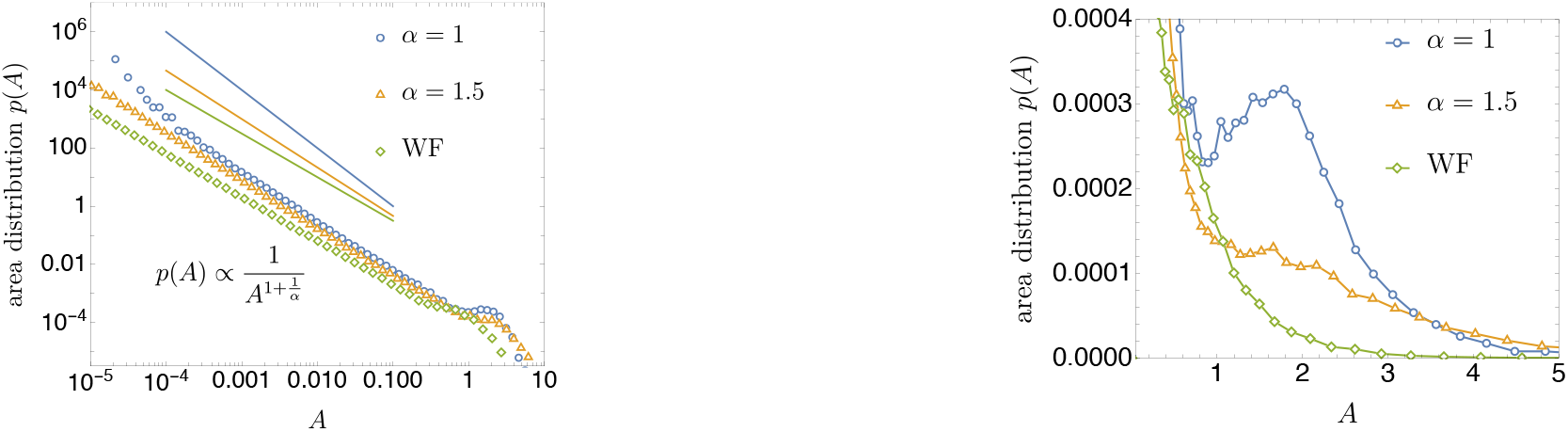
Left: The area distribution *p*(*A*) for *α* = 1, 1.5 and the Wright-Fisher model. The straight lines show the scaling-argument predictions, 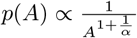. *N* = 10^6^. Right: The tail of *p*(*A*) in the large-*A* region.

Focusing on small areas, which correspond to extinct trajectories, this power-law behavior can be rationalized again from a scaling argument: First, by using Equation 3, a trajectory whose maximum frequency is *x*_*_ ≪ 1 sweeps an area roughly given by 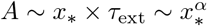 (see Figure 20), i.e., 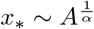. Second, from the neutrality, the cumulative probability Pr(*X*_*_ > *x*_*_) that a single mutant achieves a frequency larger than *x*_*_ before absorption is estimated as 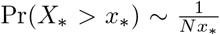. Hence, the density *p*(*x*_*_) is given by 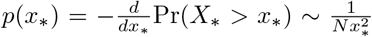. Combining these two results, we can estimate the area distribution *p*(*A*) as

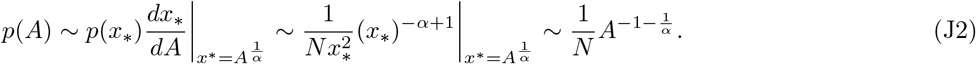

When *α* ⟶ 2 − 0 (Wright-Fisher limit), the distribution becomes 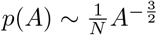, which can be analytically confirmed by solving a backward diffusion equation of the Wright-Fisher diffusion (see Appendix J-2).

The numerical results indicate that, when 1 ≤ *α* < 2, there is an uptick in the area distribution *p*(*A*), which comes from fixed trajectories (see the case of *α* = 1 in the right panel of Figure 21). The uptick becomes less pronounced as *α* increases. For the Wright-Fisher model, we can analytically prove that *p*(*A*) monotonically decreases with *A*.

## 2. Area distribution in the Wright-Fisher model

Here, we derive an analytical result of Equation J1 for the Wright-Fisher diffusion process. Consider a Langevin equation

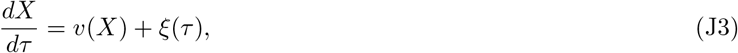

with 〈*ξ*(*τ*)*ξ*(*τ*′)〉 = 2*D*(*x*)*δ*(*τ − τ*′). Assume the initial value *X*(*τ* = 0) = *x*_0_ ∈ (0, 1) and the absorbing boundaries at *X* = 0, 1. For a given trajectory departing from *x*_0_ and ending at either one of the boundaries, we consider the “area” defined by

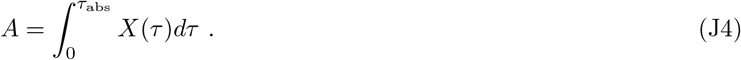

where *τ*_abs_ is the absorption time.

The area distribution Π(*A*; *x*_0_) for a given initial condition *X*(0) = *x*_0_ obeys a backward equation. To show this, we discretize the dynamics;

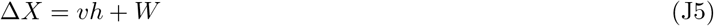

where *h* denotes a short time interval and 〈*W_i_W_j_*〉 = 2*Dhδ_i,j_*. The transition density is given by

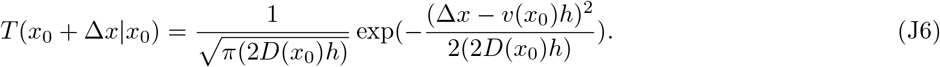

Note that

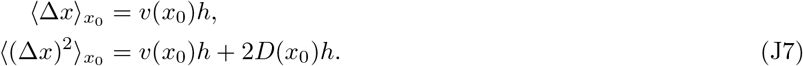

By separating a trajectory into the initial step and the remaining part, we have

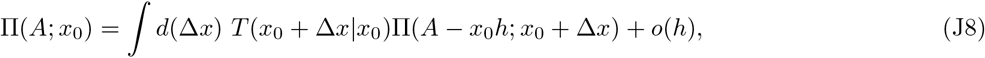

By Taylor-expanding Π(*A* − *x*_0_*h*; *x*_0_ + Δ*x*), we have

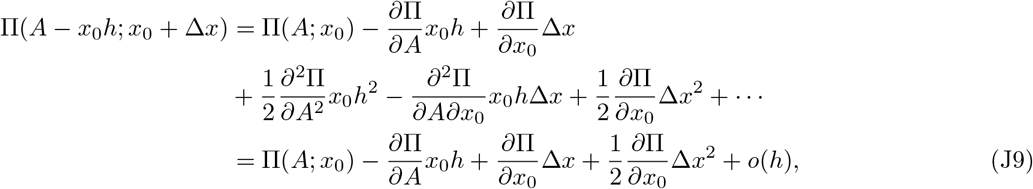

Therefore, Equation J8 becomes

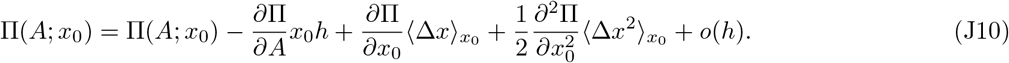

By using Equation J7, we obtain

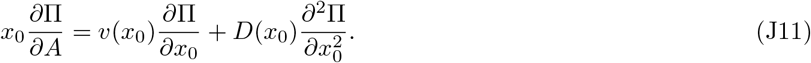

Although, in the following, we consider the area defined by Equation J4, it can be shown that, for the following integral,

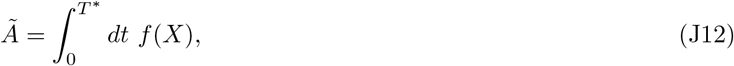

the distribution 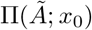 satisfies

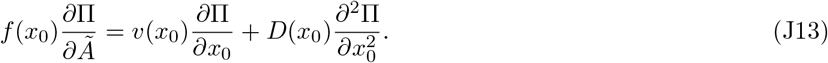

In the neutral Wright-Fisher model, *v*(*x*_0_) = 0 and *D*(*x*_0_) = *x*_0_(1 − *x*_0_). The backward equation in Equation J11 is given by

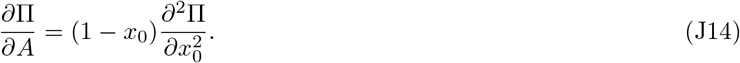

From this equation, it follows that Π(*A|x*_0_) monotonically decreases with *A*_0_ because the spectrum of the operator 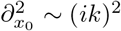 is non-positive.

We can determine the area distribution *p*(*A*) analytically at least for small *A*. We are interested in the invasion by a single mutant, 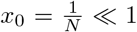. Furthermore, for the purpose of determining the behavior for small areas, we expect that we can ignore the presence of the high-frequency boundary *x* = 1 and solve the problem on the semi-infinite line *x*_0_ ∈ (0, ∞). Therefore, we consider the following problem:

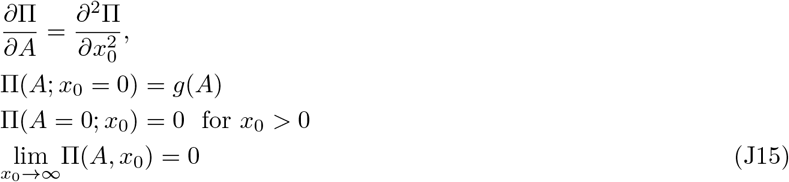

In our case, *g*(*A*) = *δ*(*A*), because the trajectory starting from *x*_0_ = 0 has *A* = 0.

For a function *f* (*A*) of *A*, we write the Laplace transformation as

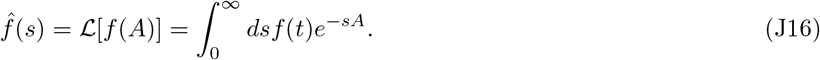

By take the Laplace transform with respect to *A*, we have

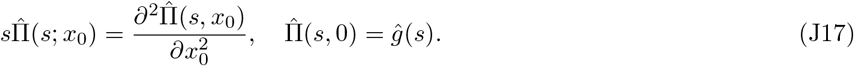

The solution is

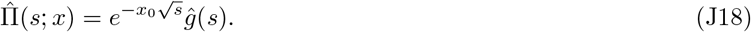

We take the inverse of the Laplace transformation,

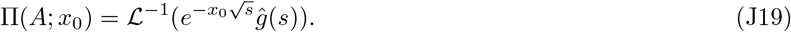

From the convolution theorem, this is given by the convolution of 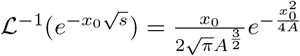 and *g*(*A*);

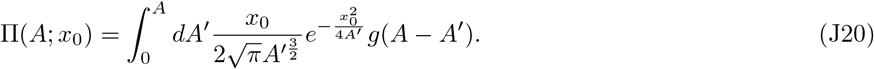

When *g*(*A*) = *δ*(*A*), we have

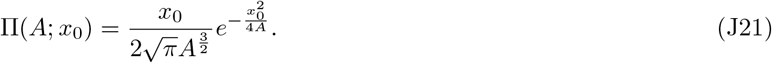

Especially, when *x*_0_ = 1*/N*, we have

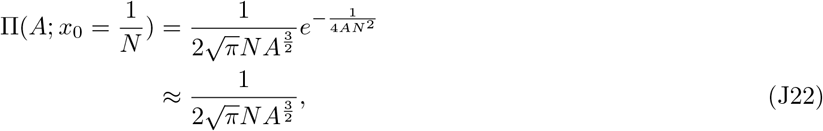

where we have used 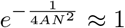 since only areas larger than 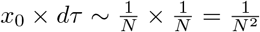 are meaningful for a finite-size population.

## Appendix K. Forward-in-time behaviors of the Eldon-Wakeley model

Here, we present simulation results of the median allele frequency and the median and mean square displacements in the Eldon-Wakeley model [11] (see also [52]). As shown below, unlike our model, these quantities do no exhibit sustained power-law behaviors, because of the existence of a characteristic size *Ψ* in the offspring distribution.

We consider the neutral Eldon-Wakeley model, where the following offspring distribution *P_U_* (*u*) is given by (see Equation (7) in [11]);

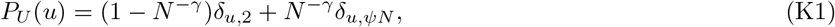

where *δ_a,b_* is the Kronecker delta. *Ψ* ∈ (0, 1) and the parameters characterizing how large and frequent ‘sweepstakes’ are.

The limiting process as *N* ⟶ ∞ depends on *γ* (see Equation (9) in [52]). For *γ* > 2, the process is the same as the Wright-Fisher diffusion, while, for *γ* < 2, it is described by a jump process whose backward-time generator 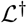 is given by

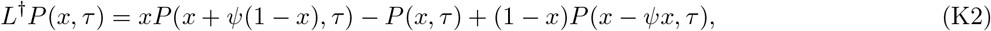

where the continuous time *τ* is related with generations *t* by *τ* = *t/N^γ^*. The first term of the generator represents a frequency-increasing jump *x ⟶ x* + *Ψ*(1 − *x*) with rate *x*, while the last one represents a frequency-decreasing jump *x ⟶ x − Ψx* with rate 1 − *x*.

Figure 22 shows numerical simulation results for the median of allele frequencies and the median/mean square displacements. The median frequency for a small initial frequency *x*_0_ ≪ 1 is well described by 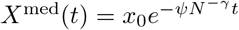 (Figure 22A). This exponential decay can be expected from the generator in Equation K2; for *x* ≪ 1, frequency-increasing jumps (with rate *x*) are unlikely to occur, and an allele frequency typically decreases by − *Ψx* with rate 1 − *x* ≈ 1. Thus, the median frequency in the Eldon-Wakeley model does not exhibit a power-law behavior.

**Figure 22.**
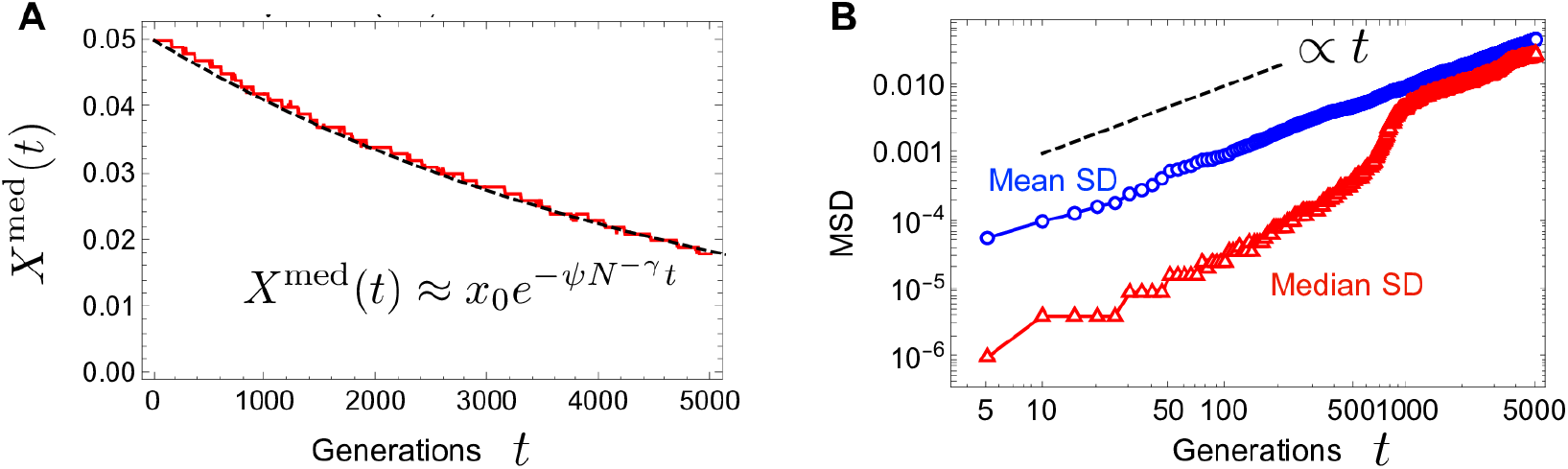
Simulation results of the Eldon-Wakeley model. (A) The median frequency of the Eldon-Wakeley model (red solid) and 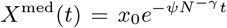 (black dashed). *N* = 10^3^, *γ* = 1, *Ψ* = 0.1, *x*_0_ = 0.05. (B) The mean and median square displacements (blue and red curves, receptively). The black dashed line 1*/t* indicates the expectation from the Wright-Fisher (or Moran) model. *N* = 10^3^, *γ* = 1, *Ψ* = 0.2, *x*_0_ = 0.5.

As for frequency fluctuations, while the mean SD exhibits a normal diffusion as in the Moran (or Wright-Fisher) model, i.e., Mean SD ∝ *t*, the median SD does not exhibit a sustained power-law behavior (Figure 22B); in a short- and long time scales, the median SD exhibits a normal diffusion (Median SD ∝ *t*), but, for an intermediate timescale (*t* ~ 500 − 1000 generations in the figure), it increases more rapidly than expected from a normal diffusion.

## Footnotes

1 *η*\* is obtained from 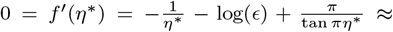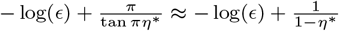.

2 Although the magnitudes of *−*1 and 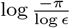 are small compared to −log *ϵ*, we need to retain these two terms because *f* (*η*^*^) contributes to *I_ϵ_* through *e^f^*^(*η*^*).

3 Note that the definition of the parameter *α_H_* in [21] is different from our definition of *α*. For 1 < α < 2, which corresponds to the semi-pushed wave region *−*1 < α_H_ < 0, the two definitions are related by *−α_H_* = *α −* 1.

4 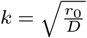 4 for 0 < B < 2, and 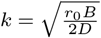 for *B ≥* 2 [21].

